# Filopodia-mediated basement membrane assembly at pre-invasive tumor boundaries

**DOI:** 10.1101/2021.10.22.464987

**Authors:** Emilia Peuhu, Guillaume Jacquemet, Colinda LGJ Scheele, Aleksi Isomursu, Ilkka Paatero, Kerstin Thol, Maria Georgiadou, Camilo Guzmán, Satu Koskinen, Asta Laiho, Laura L Elo, Pia Boström, Pauliina Hartiala, Jacco van Rheenen, Johanna Ivaska

**Author notes:** Contact information: Johanna Ivaska, Turku Bioscience Centre, University of Turku, Tykistökatu 6, FI-20520 Turku, Finland. equal contribution (Emilia Peuhu, Guillaume Jacquemet).

## Abstract

Ductal carcinoma in situ (DCIS) is a pre-invasive stage of breast cancer, where the tumor is encapsulated by a basement membrane (BM). At the invasive phase, the BM barrier is compromised enabling tumor cells to escape into the surrounding stroma. The molecular mechanisms that establish and maintain an epithelial BM barrier in vivo are poorly understood. Myosin-X (MYO10) is a filopodia-inducing motor protein implicated in metastasis and poor clinical outcome in patients with invasive breast cancer (IBC). We compared MYO10 expression in patient-matched normal breast tissue and DCIS lesions and found elevated MYO10 expression in DCIS samples, suggesting that MYO10 might facilitate the transition from DCIS to IBC. Indeed, MYO10 promoted the formation of filopodia and cell invasion in vitro and positively regulated the dissemination of individual cancer cells from IBC lesions in vivo. However, MYO10-depleted DCIS xenografts were, unexpectedly, more invasive. In these xenografts, MYO10 depletion compromised BM formation around the lesions resulting in poorly defined tumor borders and increased cancer cell dispersal into the surrounding stroma. Moreover, MYO10-depleted tumors showed increased EMT-marker-positive cells, specifically at the tumor periphery. We also observed cancer spheroids undergoing rotational motion and recruiting BM components in a filopodia-dependent manner to generate a near-continuous extracellular matrix boundary. Taken together, our data identify a protective role for MYO10 in early-stage breast cancer, where MYO10-dependent tumor cell protrusions support BM assembly at the tumor-stroma interface to limit cancer progression, and a pro-invasive role that facilitates cancer cell dissemination at later stages.

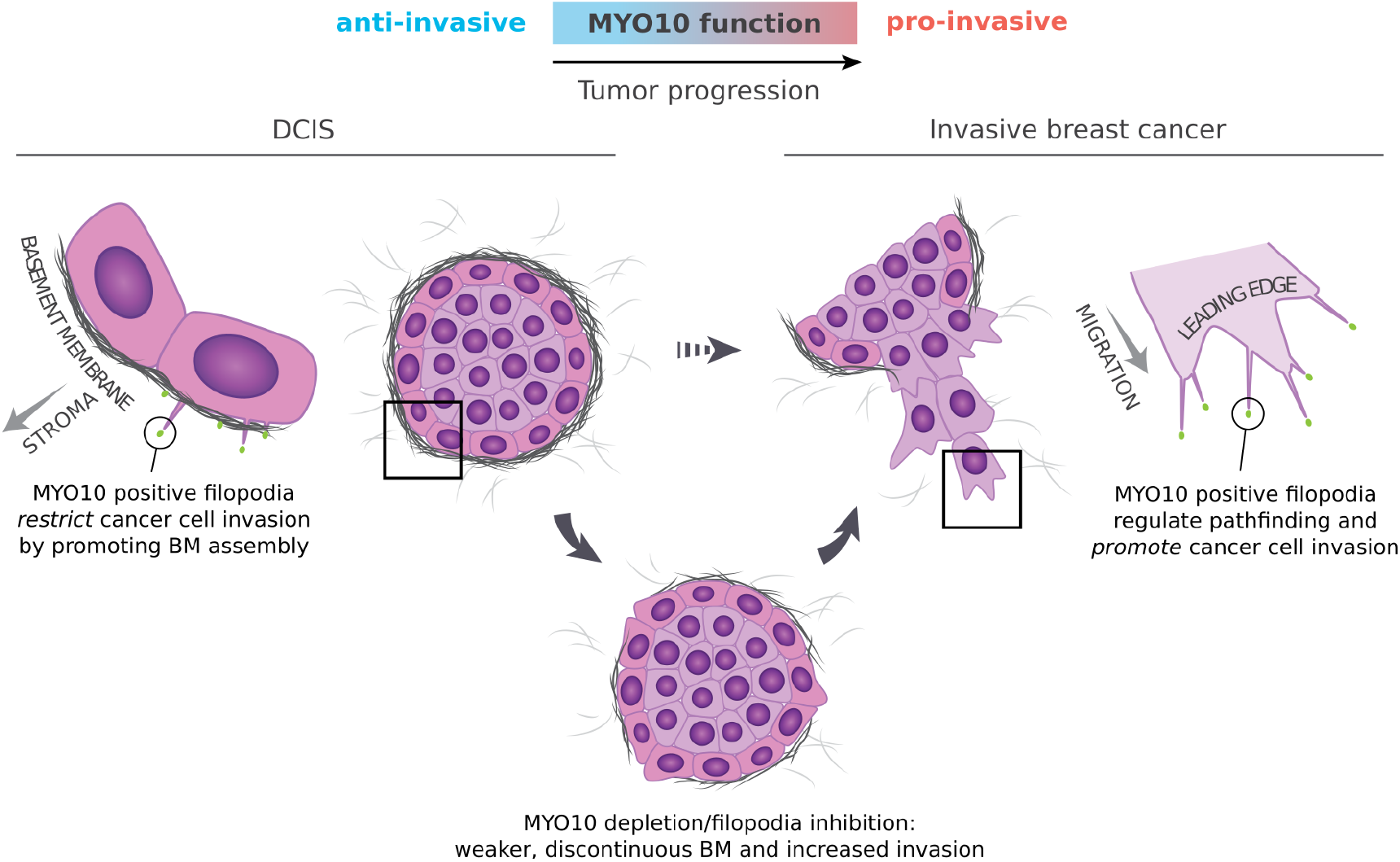

**Highlights:**

- Filopodia sculpt the tumor-proximal stroma in pre-invasive ductal carcinoma *in situ* (DCIS).
- Filopodia-dependent basement membrane (BM) assembly limits invasive transition of DCIS-like tumors *in vivo*.
- Loss of MYO10-dependent filopodia impairs BM assembly and induces an EMT-like phenotype at the tumor-stroma interface *in vivo*.
- MYO10 filopodia are anti-invasive in DCIS but facilitate dissemination in invasive breast cancer.

## Introduction

Despite recent therapeutic advances, breast cancer remains a significant cause of death among women (Ferlay et al., 2018). Breast cancer is particularly impervious to established therapies at the later stages of the disease when tumors have become invasive and metastatic. A critical step in breast cancer progression is transitioning from ductal carcinoma *in situ* (DCIS) to invasive breast cancer (IBC). DCIS is surrounded by a basement membrane (BM), acting as a barrier between the epithelial cells and the surrounding stroma. Transition to IBC involves cancer cells breaching their BM barrier and invading into the surrounding tissue, either as single cells or as a stream (Clark and Vignjevic, 2015), such that, in IBC, the BM is virtually absent. DCIS is considered a benign tumor but also a non-obligate precursor of IBC, and thus, a significant proportion of patients with DCIS diagnosis eventually progress to IBC (Betsill et al., 1978; Ryser et al., 2019). Therefore, understanding how cancer cells interact with and breach the BM is of high clinical and therapeutic interest.

BMs are thin, dense sheets of specialized extracellular matrix (ECM) molecules surrounding epithelial tissues (Yurchenco and Patton, 2009). They are composed of a three-layer ECM network, the inner layer mainly containing laminins and the outer layer formed of type IV collagen. These two layers are interlinked by several additional ECM molecules, including nidogen, lumican, and perlecan. BMs are effective biological barriers maintained by constant turnover and remodeling of ECM components (Keeley et al., 2020; Matsubayashi et al., 2020). In addition, BMs regulate epithelial architecture by establishing polarity and by providing survival cues and mechanical support.

Cancer cells can utilize specialized protrusions, such as invadopodia, to traverse BMs (Eddy et al., 2017). Invadopodia contain proteases that can degrade ECM molecules, releasing pro-invasive soluble cues and promoting the transition from *in situ* to invasive breast carcinoma (Ferrari et al., 2019; Lodillinsky et al., 2016; Monteiro et al., 2013). In addition, stromal cells may facilitate cancer cell invasion by physically remodeling BMs (Glentis et al., 2017). To date, most of the research has focused on elucidating the mechanisms by which cancer cells breach established BMs. In contrast, very little is known about how cancer progression is coupled to general BM alterations and whether cancer cells themselves could contribute to BM assembly and maintenance.

Filopodia are small and dynamic finger-like actin-rich protrusions that interact with the ECM. Filopodia contain cell-surface receptors, such as integrins, cadherins, and growth factor receptors that can interact with and interpret a wide variety of cues (Fierro-González et al., 2013; Jacquemet et al., 2019; Valenzuela and Perez, 2020). Accordingly, filopodia are essential in guiding key cellular processes such as development, angiogenesis, immune surveillance, and wound healing (Jacquemet et al., 2015). Filopodia are also associated with increased invasion and metastasis in several cancer types and support of cancer cell survival at metastatic sites (Jacquemet et al., 2017; Shibue et al., 2012, 2013).

Filopodia are relatively well-defined structures in cells on 2D substrates (Jacquemet et al., 2019), and filopodia-like protrusions have also been demonstrated in 3D substrates and *in vivo* (for instance, (Jacquemet et al., 2013; Liu et al., 2018; Millard and Martin, 2008). However, it is often unclear how these structures compare to the 2D filopodia, and how they contribute to cancer cell invasion. Furthermore, much of the research on cell protrusions in cancer has focused on cancer cell-intrinsic properties facilitating migration, invasion, and navigation through the complex 3D stroma. The possibility of cell protrusions regulating different aspects of the tumor microenvironment during cancer progression, on the other hand, is not well understood.

MYO10 is a well-established inducer of filopodia in many cell types and a positive regulator of migration and invasion of single and collectively migrating cells (Arjonen et al., 2014; Berg and Cheney, 2002; Summerbell et al., 2020). In IBC, MYO10 expression is upregulated by mutant p53, leads to metastasis, and correlates with poor patient outcomes (Arjonen et al., 2014; Cao et al., 2014).

To understand whether MYO10 filopodia contribute to the DCIS-to-IBC transition, we interrogated MYO10 expression in patient samples. MYO10 expression was increased in DCIS compared to the normal mammary epithelium. Rather surprisingly, the depletion of MYO10 expression intensified the loss of DCIS-like morphology in a xenograft model of breast cancer progression. It also induced an EMT-like phenotype at the tumor borders and compromised BM formation around the DCIS lesions, leading to the dispersal of carcinoma cells into the surrounding stroma. Furthermore, imaging experiments demonstrated that MYO10-induced protrusions contribute to ECM assembly. Taken together, the data reported here support a model where MYO10 plays a dual role in breast cancer. At the DCIS stage, MYO10 facilitates cancer ECM assembly and the generation of a tumor-stroma barrier, whereas, at the IBC stage, MYO10 induces cancer cell migration and dissemination.

## Results

### Filopodia depletion limits collective cell migration in vitro and individual cancer cell motility in vivo

The evolution of cancer is frequently modeled using related cell lines with progressing malignancy. One such model is the widely used immortalized breast epithelial cells (MCF10A), their H-Ras transformed variants (MCF10AT cell line) that are tumorigenic as xenografts, and the tumorigenic and invasive cells derived from MCF10AT (MCF10DCIS.COM cell line; (Dawson et al., 1996; Miller et al., 2000)). Xenografts of the MCF10DCIS.COM cells recapitulate the different stages of DCIS progression mimicking human disease (Miller et al., 2000; Behbod et al., 2009; Lodillinsky et al., 2016). Previously, we have shown that MCF10DCIS.COM cells display high numbers of filopodia as they invade collectively *in vitro* (Jacquemet et al., 2017). However, the role of MYO10 in these cells has remained unknown. To investigate MYO10 dependency in the generation of these filopodia, we silenced MYO10 expression using two independent short hairpin RNA (shRNA). Both shRNAs silenced MYO10 efficiently and reduced filopodia density to a similar degree (**Fig. S1A-C**). Subsequently, we pooled four MYO10-depleted single-cell clones to create the shMYO10 DCIS.com cell line used in this study (**Fig. 1A** and **Fig. S1D-E**). Silencing of MYO10 led to a marked reduction in filopodia density and length in cells migrating in 2D (**Fig. 1B**) and decreased cell invasion speed through collagen in a 2D overlay assay (**Fig. 1C** and **Video 1**). Furthermore, high-resolution live-cell imaging revealed that in the absence of filopodia, shMYO10 DCIS.com cells switch to a lamellipodia-driven mode of collective cell migration (**Fig. 1D** and **Video 2**). Taken together, these data indicate that MYO10 regulates the protrusive activity and collective migration of DCIS.com cells *in vitro*.

**Figure 1:**
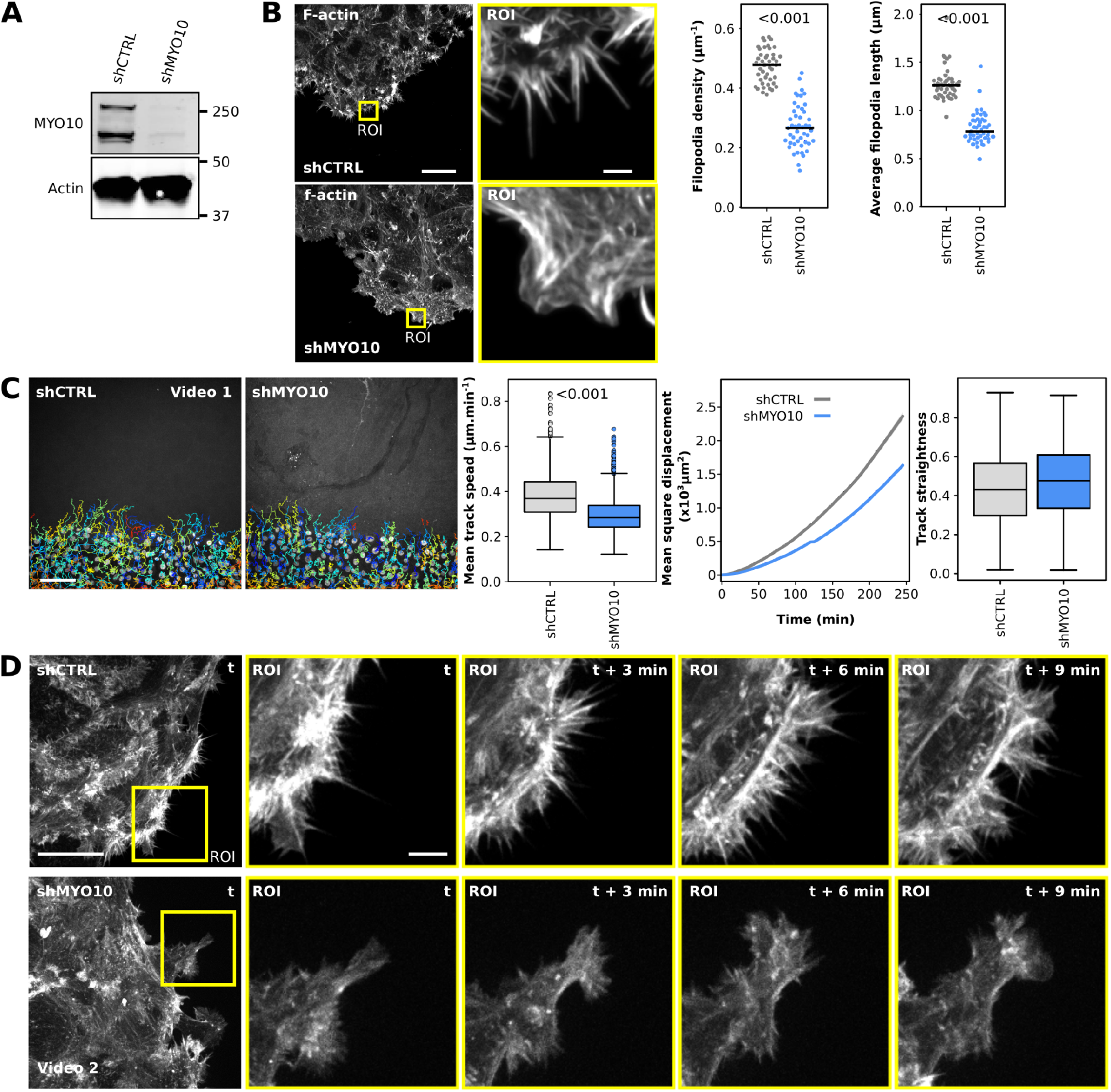
Filopodia depletion triggers a switch to lamellipodia-driven migration and limits cell motility. (**A**): shCTRL and shMYO10 DCIS.com cells were lysed, and MYO10 protein levels were analyzed by western blot. (**B**): shCTRL and shMYO10 DCIS.com cells were left to migrate underneath a collagen gel for two days (d), fixed, stained, and imaged using a spinning-disk confocal microscope. A representative field of view is displayed. Yellow squares highlight regions of interest (ROIs) that are magnified. Scale bars: (main) 25 μm; (inset) 2 μm. Filopodia density and the average filopodia length were analyzed using FiloQuant. Results are displayed as dot plots (n > 45 fields of view analyzed per condition; three independent experiments; randomization test). (**C**): shCTRL and shMYO10 DCIS.com cells were left to migrate underneath a collagen gel for 1 d, incubated with SiR-DNA (to visualize nuclei), and imaged live using a spinning-disk confocal microscope (20x air objective). Cells were then automatically tracked using StarDist and TrackMate. A representative field of view with cell tracks is displayed (See also Video 1). Mean track speed, mean square displacement, and track straightness were calculated using the motility lab website (three independent experiments, 30 fields of view per condition, and n > 2300 cell tracks; randomization test). Scale bar: 100 μm. (**D**): shCTRL and shMYO10 DCIS.com cells were left to migrate into a collagen gel for 1 d and imaged live using a spinning-disk confocal microscope (100x objective). A representative field of view with selected time points is displayed (See also Video 2). Scale bars: (main) 25 μm; (inset) 5 μm.

### Filopodia contribute to the multicellular organization of cancer cells *in vitro* and *in vivo*

To study the role of MYO10 filopodia in organizing cells within a monolayer, we mixed shCTRL and shMYO10 cells (alternating GFP labeling of the cell lines) (**Fig. 2A**) and recorded cell migration live (**Fig. 2B** and **Video 3**). Notably, shMYO10 cells consistently lagged behind shCTRL cells, which preferentially localized to the front of the collectively migrating cells (**Fig. 2B-C** and **Video 3**). This result demonstrates that MYO10 expression segregates cancer cells within an actively migrating monolayer.

**Figure 2:**
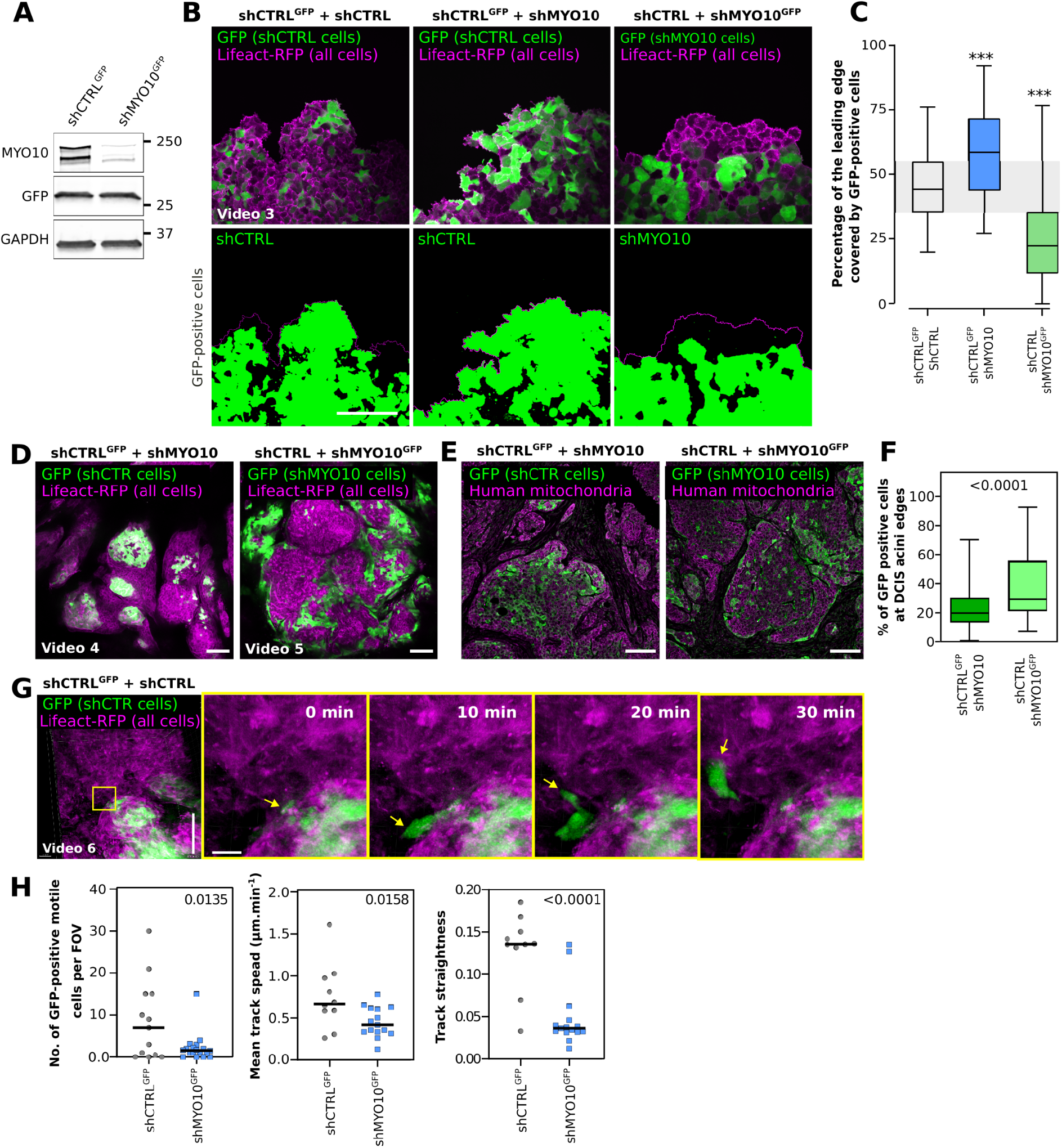
Cell competition experiments reveal that MYO10 filopodia contribute to the multicellular organization of cancer cells in vitro and in vivo. (**A**): shCTRL and shMYO10 DCIS.com cells were infected with GFP-containing lentiviruses, lysed, and their MYO10 and GFP expression levels were analyzed by western blots. A representative western blot is displayed. (**B-C**): Various shCTRL and shMYO10 DCIS.com cell lines were mixed in different combinations so that one of the cell lines is always GFP positive, and cell migration was recorded live on a spinning-disk confocal microscope (20x). Representative images are displayed. Scale bar: 200 μm. For each condition, the percentage of the leading edge covered by GFP-positive cells was measured using Fiji. (**C**): The results are displayed as Tukey box plots (n > 5266 fields of view analyzed per condition; 3 biological repeats; *** p-value < 0.001, randomization test). (**D-F**): shCTRLGFP + shMYO10 or shMYO10GFP + shCTRL DCIS.com cells were xenografted in immunocompromised mice in 1:1 ratio. The resulting xenografts were imaged by intravital tile scan imaging (n = 2) (**D**) or dissected, sectioned, and imaged using a spinning-disk confocal microscope (20x objective) (**E**). The percentage of GFP-positive cells at the edge of DCIS acini was then scored using Fiji, and results displayed as Tukey box plots (**F**) (n = 4 tumors per condition; fields of view analyzed: shCTRLGFP + shMYO10, 114; shMYO10GFP + shCTRL,103; Mann-Whitney test). Scale bars: 100 µm. (**G**): shCTRLGFP or shMYO10GFP DCIS.com cells were injected along with non-GFP cells in immunocompromised mice. Intravital imaging of tumors was conducted 25-35 days post-tumor inoculation (see also Video 6). Scale bars: (main) 100 μm (inset) 15 μm. (**H**): All visibly motile GFP-positive cells were tracked manually in three dimensions using the Imaris software, and the number of motile cells per field of view and mean track speed and track straightness (displacement/length of track) per cell were quantified (n = 4 shCTRL-GFP tumors with 2-4 FOVs per tumor; and n = 6 shMYO10-GFP tumors with 1-5 FOVs per tumor; t-test). FOV: field of view.

Cell migration also contributes to the multicellular organization within tumors (Waclaw et al., 2015). To investigate whether filopodia would also contribute to the cellular localization pattern in DCIS-like tumors *in vivo*, we used two-photon intravital microscopy (**Fig. 2D**) and tumor histology (**Fig. 2E**). We examined the localization of shCTRL and shMYO10 cells within the same tumor (from mice grafted with a 50:50 mixture of shCTRL and shMYO10 cells with GFP expressed in either shCTRL or shMYO10 cells). Strikingly, we observed obvious preferential segregation of shMYO10 cells to the edges of the tumor acini while shCTRL cells tend to accumulate toward the center (**Fig. 2D-F** and **Video 4-5**). This result indicates that MYO10 also contributes to the multicellular organization of tumor cells *in vivo*, but, surprisingly, the segregation of MYO10 depleted cells is inverted compared to the *in vitro* freely migrating setup described above (**Fig. 2B-C**). These data indicate that MYO10 may contribute to distinct cell behavior in pre-invasive confined and invasive actively migratory conditions.

Despite the preferential recruitment of MYO10 depleted cells at the edges of the acini, GFP-based single-cell tracking in tumors from live two-photon intravital imaging (**Fig. 2G**) (see Methods for details) indicated that shCTRL cells had increased single-cell migration and invasion out of the DCIS-like acini *in vivo* compared to shMYO10 cells (**Fig. 2H** and **Video 6**). This result agrees with our *in vitro* migration experiments indicating that MYO10 depletion reduces cell migration (**Fig. 1C**). Thus, MYO10 filopodia contribute to the migratory behavior and the multicellular organization of cancer cells *in vitro* and *in vivo*.

### MYO10 depletion induces cancer cells dispersal and EMT, specifically at the tumor border

These data led us to the unexpected hypothesis that MYO10-dependent filopodia may have distinct roles in early-stage breast cancer and in IBC, where MYO10 contributes to systemic metastasis (Arjonen et al., 2014; Cao et al., 2014). MYO10 is highly expressed in a subset of breast carcinomas with poor prognosis (Arjonen et al., 2014). However, MYO10 expression at the early, non-invasive stage of breast cancer has not been investigated. We obtained freshly operated tissues from a set of patients diagnosed with DCIS, a pre-invasive and BM-confined tumor. MYO10 mRNA expression, detected using RNAscope in-situ hybridization, was significantly higher in the DCIS lesions than in the normal tissue and overall MYO10 was expressed at a very low level in the normal epithelium **(Fig. 3A and Fig. S2A-B**). This indicates that MYO10 may play a role already in the early stages of tumor progression.

**Figure 3:**
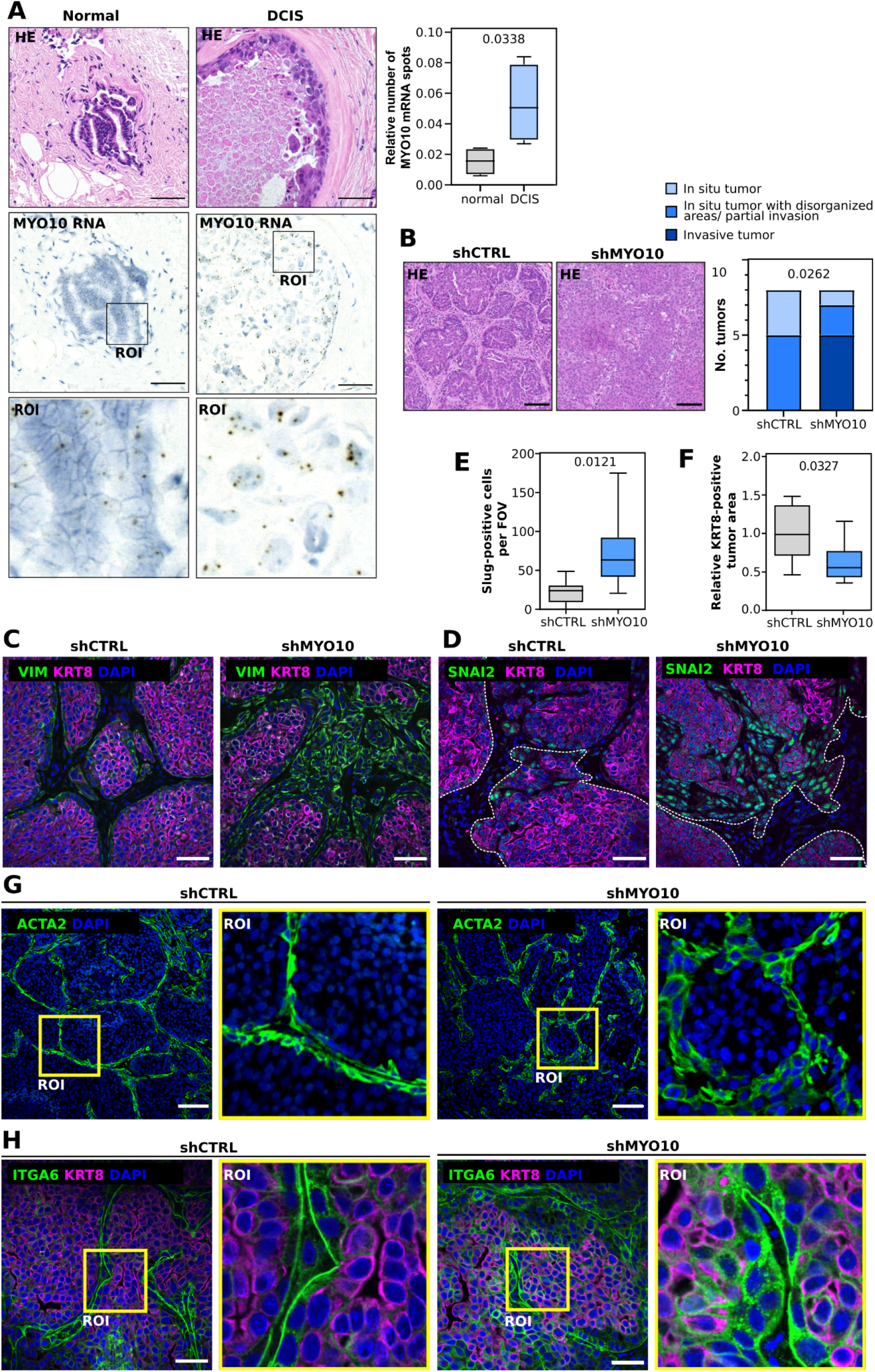
*In vivo* MYO10 depletion induces cancer cells dispersal and EMT, specifically at the tumor border. (**A**): *In situ* labeling of MYO10 mRNA in normal and DCIS regions of a human breast sample (images representative of 4 patient samples per condition). MYO10 mRNA can be visualized by the dots visible in the magnified ROI. Scale bars: (main) 100 μm; (inset) 20 μm. Representative images of and quantification of the number of MYO10 mRNA spots relative to normal/DCIS tissue area are shown (n = 4 patient samples, Student’s t-test). (**B**): shCTRL and shMYO10 cells were injected subcutaneously in NOD.scid mice. At 25 days post-injections, tumors were dissected, and tissue sections were stained as indicated and imaged. Representative images of tumor histology and quantification of invasiveness are shown (n = 8 tumors, Chi-square test). (**C-D**): Representative images of day 25 tumor sections labeled for vimentin (human-specific antibody, VIM) and keratin-8 (KRT8) (**C**) or Slug (SNAI2) and KRT8 (**D**) were taken using a confocal microscope. The area of cancer cells is indicated by dashed lines in **D**. The average number of Slug-positive cells per field of view (**E**), or the relative KRT8-positive tumor area (**F**) were quantified (n=8 tumors from 2 independent experiments; unpaired t-test). (**G-H**): Representative images of tumor sections labeled for αSMA (ACTA2; **G**; n=5 tumors) or ITGA6 and KRT8 (**H**; n^shCTRL^ = 3; n^shMYO10^ = 2 tumors). Squares represent ROIs that are magnified.

To test the outcome of MYO10 depletion in vivo, we generated shCTRL and shMYO10 xenografts. The transition from DCIS to IBC in this model progresses over time in vivo (**Fig. S3A**). MYO10 silencing did not affect cell growth in 2D culture (**Fig. S3B**), and the sizes of shCTRL and shMYO10 xenograft tumors were comparable (**Fig. S3C-D**). Next, we compared the onset of invasion in shCTRL and shMYO10 xenografts by blind scoring the degree of invasion based on the tumor histology (**Fig. 3B and Fig. S3E**). As expected, at 25 days post-inoculation, shCTRL tumors were composed of DCIS-like acini or acini exhibiting partial invasion (**Fig. 3B**). In contrast, most shMYO10 tumors displayed partial or complete invasion and loss of the in situ tumor organization (**Fig. 3B**).

Breast cancer is a heterogeneous disease and presents with a varying degree of epithelial to mesenchymal transition that has been associated with increased tumor invasion. Immunostaining of the xenografts with markers for epithelial/luminal-like (keratin 8; KRT8) and mesenchymal/basal-like breast cancer cells (vimentin; VIM and the transcription factor Slug; SNAI2) revealed a notable presence of cells with mesenchymal traits, particularly at the perimeter of MYO10-depleted xenografts (**Fig. 3C-F**). Additional basal markers, α6 integrin and α smooth muscle actin (αSMA/ACTA2) that were clearly expressed at the edges of shCTRL and shMYO10 tumors, further validated the observation that basal-like cells were distributed over a broader margin at the edges of shMYO10 tumor acini (**Fig. 3G-H**). Thus, MYO10 depletion leads to an increased presence of tumor cells with mesenchymal/basal-like features that may be the cause or the consequence of the faster onset of tumor invasion in MYO10-depleted tumors.

### MYO10 promotes basement membrane assembly *in vivo*

Breaching the BM barrier is a critical step in the DCIS transition to IBC. To visualize possible BM alterations, we stained for BM components in the tumor xenografts (**Fig. 4A-D**). Tumors formed in 25 days by shCTRL cells were surrounded by clear, seemingly continuous collagen IV-positive BM. In contrast, the BM around shMYO10 tumors were harder to detect and exhibited decreased collagen IV staining (**Fig. 4A-C**). BMs contribute to the assembly of other ECM scaffolds, including the proper assembly of fibronectin (FN) fibrils (Lu et al., 2020). In line with this notion, we observed that the FN matrices in shMYO10 tumors were less developed and the ECM surrounding the tumors contained less FN than the shCTRL tumors (**Fig. 4B-C**).

**Figure 4:**
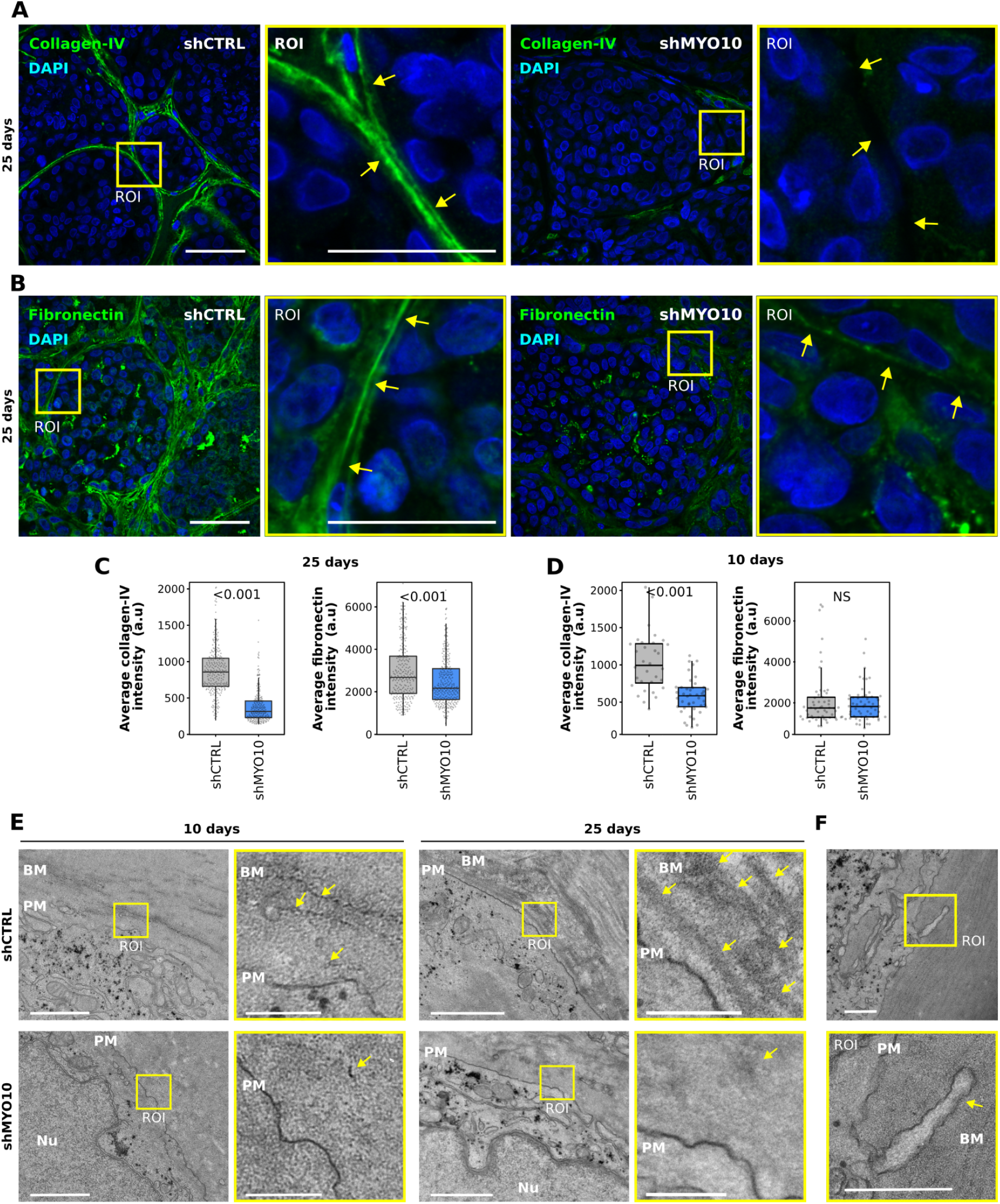
MYO10 promotes basement membrane assembly *in vivo*. (**A-D**): Tissue section of shCTRL and shMYO10 DCIS-like xenografts at day 25 (**A-C**) and day 10 (**D**) were stained for DAPI and collagen IV or fibronectin and imaged using a spinning-disk confocal microscope (63x objective). (**A-B**): Representative fields of view of day 25 DCIS-like xenografts are displayed. Scale bars: (main) 50 μm; (inset) 25 μm. (**C-D**): The average integrated density of collagen IV and fibronectin staining around the DCIS acini was measured using Fiji. The results are displayed as box plots (Day 25 xenografts, n > 233 DCIS acini from 5 tumors per condition; Day 10 xenografts, n > 32 DCIS acini from 4 tumors per condition; randomization test). **(E):** Day 10 and day 25 shCTRL and shMYO10 DCIS-like xenografts were imaged using electron microscopy to visualize the BM surrounding the DCIS acini. Representative fields of view are displayed (25-day-old xenografts, three biological repeats; ten-day-old xenografts, four biological repeats). Scale bars: (main) 1 μm; (inset) 250 nm. **(F):** Day 25 shCTRL DCIS-like xenografts were imaged using electron microscopy to visualize the protrusions surrounding the DCIS acini. Scale bars: (main) 500 nm; (inset) 500 nm. For all panels, p-values were determined using a randomization test. NS indicates no statistical difference between the mean values of the highlighted condition and the control. PM: plasma membrane; Nu: nucleus.

A similar defect in collagen IV deposition around shMYO10 acini was apparent at an earlier stage of tumor development when the acini form (10 days post-inoculation) (**Fig. 4C**). While FN staining intensity appeared equal in shCTRL and shMYO10 tumors, visible FN fibrils, constituting a continuous network, were mostly absent in shMYO10 acini (**Fig. 4D, Fig. S4**). Instead, bright FN puncta, which are reminiscent of folded FN not yet assembled into filaments, could be observed at the edges of shMYO10 acini.

The lack of BM assembly in the MYO10-depleted tumors was also observed with electron microscopy (EM). In negatively-stained EM, BMs were visible (as dark fibers) at the edges of shCTRL acini but could not be easily observed in shMYO10 tumors (**Fig. 4E**). Moreover, occasional filopodia-like protrusions were observed at the edge of the shCTRL, but not in shMYO10 DCIS-like xenografts (**Fig. 4F**). Taken together, these data indicate that the BM of shMYO10 acini is already defective while the tumors are forming and point to inadequate BM production or assembly rather than degradation, which typically occurs at a much later stage in tumor progression (Lodillinsky et al., 2016).

### ECM production is up-regulated in MYO10-depleted tumors

The tumor microenvironment is composed of ECM generated by stromal cells and tumor cells. However, in human clinical samples or genetically engineered mouse tumor models, the distinction of the ECM-producing cells is complicated. We took advantage of our ability to distinguish gene expression changes in the tumor (human genes) and the stroma (mouse genes) in our model system (see methods for details) and performed mRNA sequencing of shCTRL and shMYO10 tumors (at 25 days post-inoculation) to investigate the expression of BM components. MYO10 expression was decreased in shMYO10 tumors, validating our approach (**Fig. S5A**). Overall, stromal gene expression profiles were nearly identical in shCTRL and shMYO10 tumors (**Fig. 5A**). In contrast, gene expression profiles of shCTRL and shMYO10 tumors were distinct, with many differentially expressed genes detected between the tumors (**Fig. 5B** and **table S1**). Expression of the filopodia-inducing proteins FMNL3 and neurofascin were increased in shMYO10 tumors, indicating a possible compensatory mechanism to promote filopodia formation when MYO10 is downregulated (**Fig. S5B-C**).

**Figure 5:**
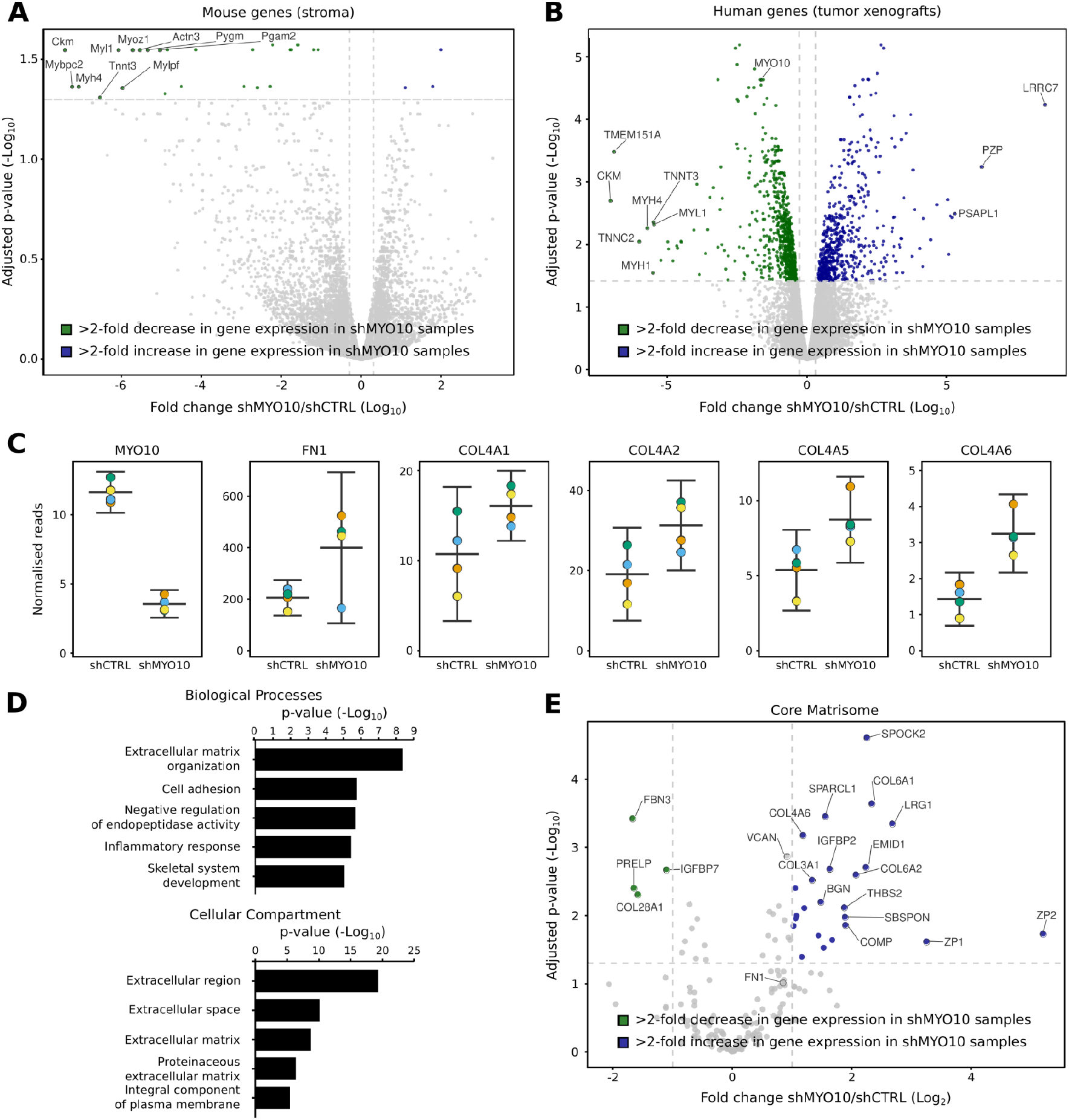
MYO10 depletion drives the expression of ECM genes by cancer cells. (**A-E**): 25-day-old shCTRL and shMYO10 DCIS-like xenografts were dissected and their RNA sequenced. The expression levels of the mouse genes (tumor stroma) and of the human genes (tumor) were analyzed separately (Table S1, see methods for details, four different mice per condition). (**A-B**) The overall gene expression changes in the stroma (**A**) or the tumors (**B**) upon MYO10 silencing are displayed as volcano plots. Genes with at least a two-fold increase in their expression levels upon MYO10 silencing are highlighted in blue, while genes with at least a two-fold decrease in their expression levels upon MYO10 silencing are highlighted in green. The most affected genes and MYO10 are annotated. (**C**): The expression levels of MYO10 and selected ECM genes FN1, COL4A1, COL4A2, COL4A5, and COL4A6 in 25-day-old shCTRL and shMYO10 DCIS-like xenografts were measured by RNAseq and displayed as SuperPlots. (**D**): Gene ontology-based functional annotation analyses of human genes overexpressed in shMYO10 tumors (Biological Process and Cellular Compartment) were performed using DAVID (Huang et al., 2009). The top five categories (based on their adjusted p-values) are displayed. (**E**): Volcano plots highlighting the changes in core Matrisome gene expression upon MYO10 silencing (as defined by the Matrisome project, http://matrisomeproject.mit.edu/).

Expression of several ECM molecules, including collagen IV (COL4A1, COL4A2, COL4A5, and COL4A6), collagen VI (COL6A1), laminin (LAMA1), and fibronectin (FN1), were increased in shMYO10 tumors (**Fig. 5C** and **Fig. S5C**), which was unexpected considering the inadequate BM generation in these tumors (**Fig. 4**). Furthermore, gene ontology analyses and annotation of our dataset using the Matrisome database (Naba et al., 2012, 2016, 2017) confirmed that MYO10-depleted xenografts produce more ECM proteins overall (**Fig. 5D** and **5E**). Taken together, these data indicate that the tumor cells are a significant source of the BM components in these DCIS-like tumors and that MYO10 depletion does not reduce BM component production but instead leads to an overall increase in ECM production *in vivo*.

### Filopodia protrusions engage ECM components in 3D

To assess whether filopodia contribute to ECM assembly, we set up 3D spheroid assays, reconstituting BM component assembly *in vitro*. Previous work using non-transformed mammary epithelial cells indicates that 3D matrigel-embedded spheroids undergo rotational motion to assemble BM (Wang et al., 2013). We first investigated the ability of DCIS.com cells to rotate in 3D cultures. Around half of the structures rotated (**Fig. 6A and Video 7**), indicating these cells may have the ability to weave exogenous ECM around the spheroids. We tested this by incubating the spheroids with fluorescently labeled ECM molecules and observed a clear layer of ECM recruited around the spheroids, immediately adjacent to the outermost cells with filopodia protruding through it (**Fig. 6B**). This was also seen while imaging spheroids live, indicating that this is not a fixation artifact (**Video 8**).

**Figure 6:**
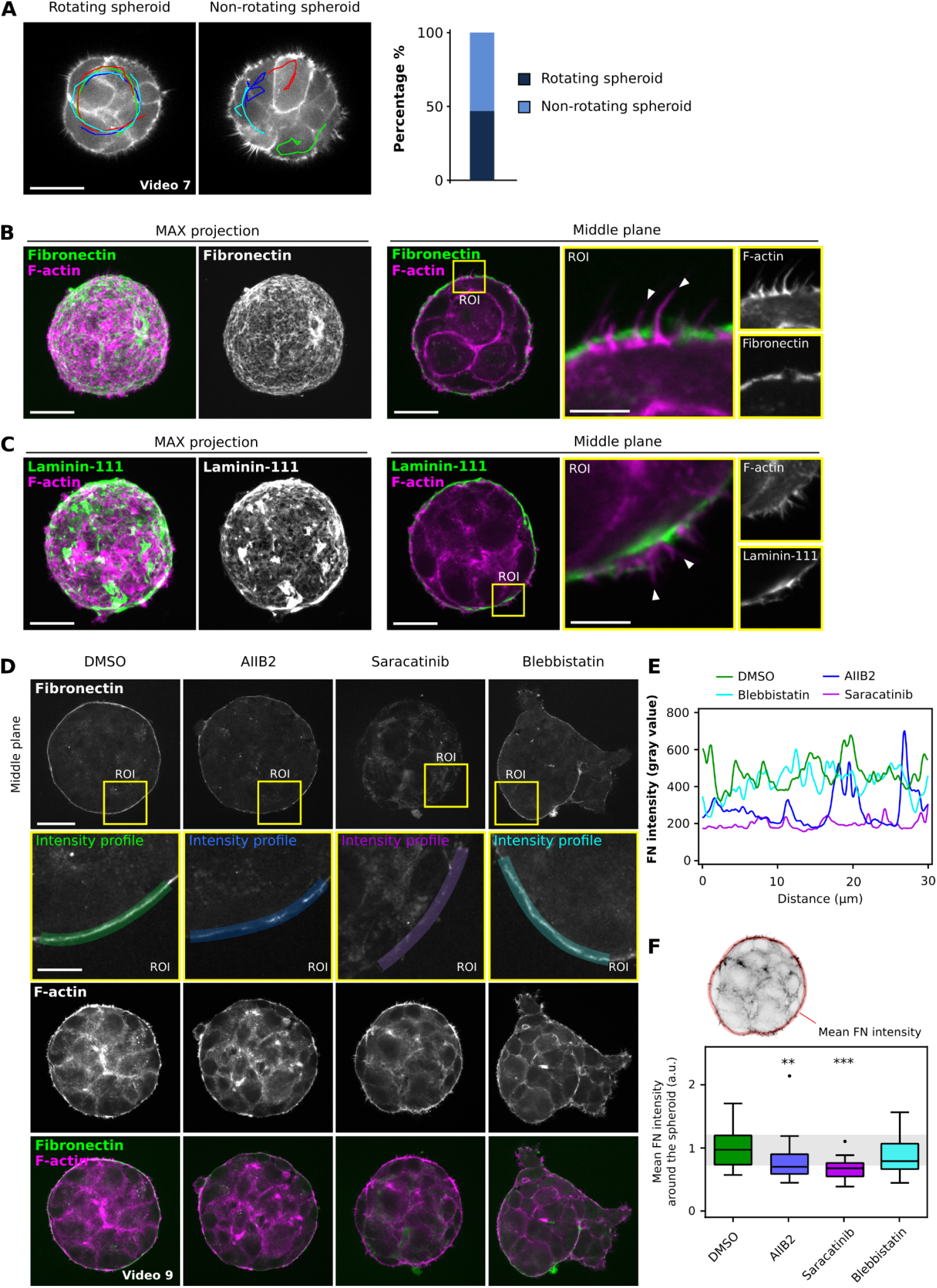
Filopodia protrusions engage ECM components in 3D. (**A**): To highlight cellular behavior within spheroids, migration tracks (red, light, dark blue, and green) are displayed on top of two different spheroids. In addition, the proportion of rotating spheroids over non-rotating spheroids is displayed. Results are from three independent experiments (81 movies analyzed). (**B-C**): Lifeact-mRFP-expressing DCIS.com cells were seeded as single cells in Matrigel. They were allowed to form spheroids for three days in the presence of fluorescently labeled FN (**B**) or laminin-111 (**C**). Samples were fixed and imaged using a spinning-disk confocal microscope. Representative fields of view highlighting the spheroids’ middle and maximum intensity projections are displayed. Scale bars: (main) 20 μm; (inset) 5 μm. (**D-E**): Fluorescence images (**D**) and intensity profiles (**E**) depicting the accumulation of exogenous fibronectin on spheroids treated with integrin/actomyosin-targeting compounds. Lifeact-mRFP-expressing DCIS.com cells were allowed to form spheroids for two days before the cultures were supplemented with fluorescently labeled FN and DMSO, blebbistatin, integrin β1-blocking antibody (AIIB2), or saracatinib. After an overnight treatment, the samples were fixed and imaged using a spinning-disk confocal microscope. Scale bars: (main) 30 μm; (inset) 10 μm. (**F**) Mean concentration of accumulated FN around each spheroid. Results are displayed as Tukey box plots (n = 23-32 spheroids; 2 biological repeats; **p-value = 0.0097, ***p-value < 0.001, Kruskal-Wallis one-way ANOVA with Dunn’s posthoc test).

Filopodia tips have small integrin-containing adhesions, which interact with the ECM to facilitate filopodia stability (Jacquemet et al., 2016; Miihkinen et al., 2021). Downstream of integrins, filopodia are dependent on Src-kinase activity, whereas inhibition of cell body contractility with Myosin-II inhibitor blebbistatin triggers elongation of filopodia (Jacquemet et al., 2016; Stubb et al., 2020). We observed that relatively short overnight incubation with exogenous ECM components was sufficient for established spheroids to enwrap in labeled ECM (**Fig. 6C and Video 9**). This enabled us to test filopodia modulators without interfering with 3D spheroid generation. Inhibiting a subfamily of integrins with an integrin β1 blocking antibody (AIIB2) reduced FN recruitment to spheroids significantly (**Fig. 6C**). The effect of Src inhibition was even more pronounced, resulting in a weaker FN signal around the spheroids and displacement of the signal to the spheroid interior (**Fig. 6C and Video 9**). The BM barrier became discontinuous in both cases compared to the FN ECM recruited by control-treated spheroids (**Fig. 6D-E**). In stark contrast, blebbistatin triggered a prominent invasion of collective, filopodia-rich cell strands from the spheroids (**Fig. 6C**). The BM barrier around the non-invading areas of the spheroids appeared intact but was clearly displaced at the invasive areas. Finally, we performed the same ECM assembly assay in MYO10-depleted spheroids. In line with the previous results, we observed that silencing of MYO10 decreased filopodia density and resulted in a much weaker accumulation of ECM molecules around the spheroids (**Fig. 7A-B**), indicating that MYO10-induced filopodia contribute to ECM assembly in 3D. Overall, these 3D ECM recruitment assays suggest a functional role for filopodia in ECM assembly.

**Figure 7:**
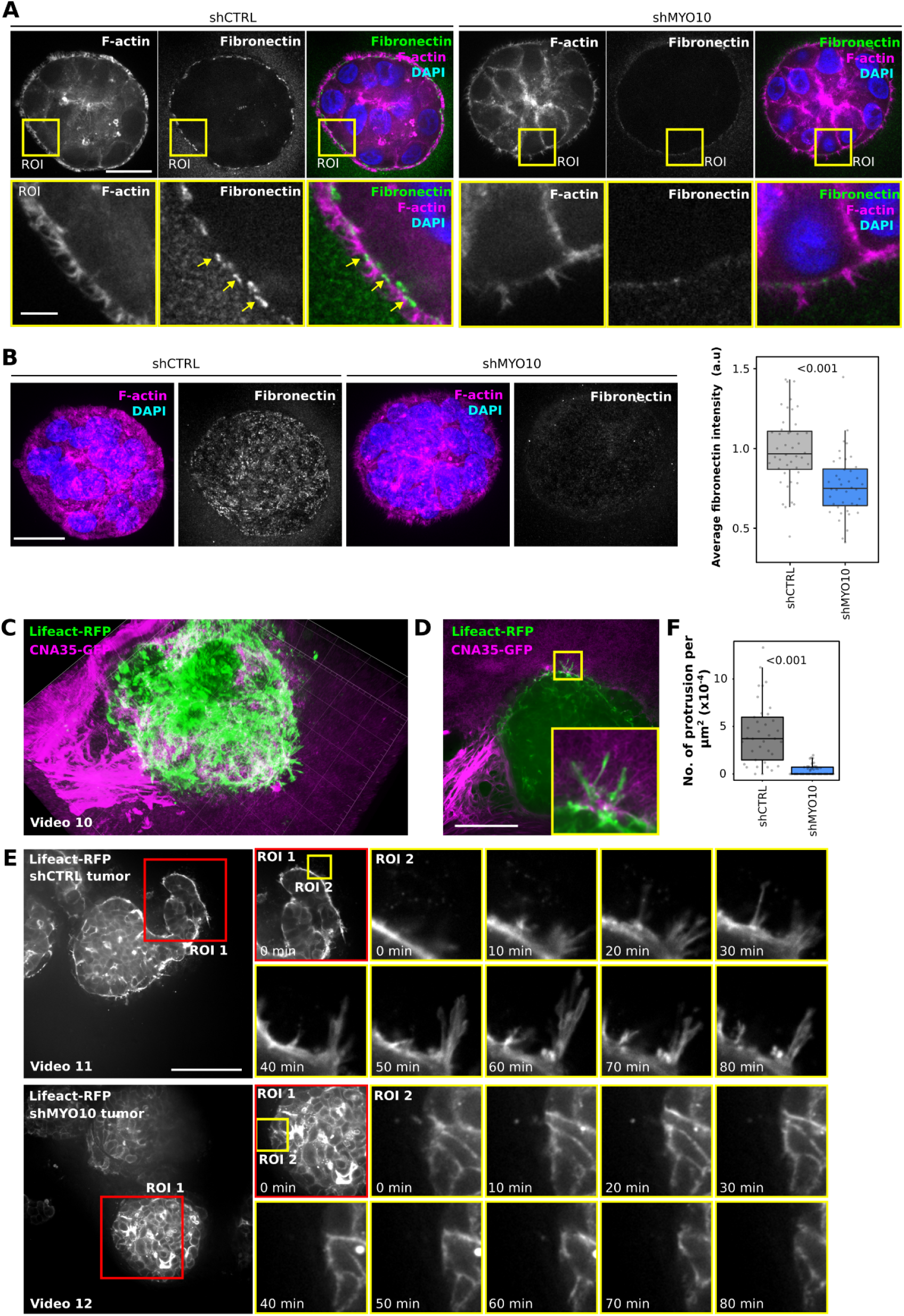
MYO10 modulates ECM assembly *in vitro* and cell protrusions at the tumor boundary ex vivo. (**A-B**): shCTRL and shMYO10 DCIS.com cells were allowed to form spheroids as in (**Fig 6**) in the presence of fluorescently labeled fibronectin and imaged using a spinning-disk confocal microscope (63x objective). Representative fields of view highlight the spheroids’ middle planes (**A**) and SUM projections (**B**). Scale bars: (main) 25 μm; (inset) 5 μm. From the SUM projections, the average fibronectin intensity per spheroid was quantified using Fiji (n = 4 biological repeats; shCTRL, n = 44 spheroids; shMYO10, n = 40 spheroids; randomization test). Yellow squares indicate ROIs that are magnified. (**C-F**): Twenty-five-day-old DCIS-like xenografts were imaged live ex-vivo using a spinning-disk confocal microscope (40x objective, ORCA camera). Scale bars: (main) 100 μm. (**C-D**): shCTRL xenografts were incubated with the fibrillar collagen probe CNA35-GFP before imaging. A 3D reconstruction (**C**, see also Video 10) and a single Z plane (**D**) of a representative field of view is displayed. (**E-F**): shCTRL and shMYO10 xenografts were imaged live over an extended time. (**E**): Representative fields of view are shown (see also Videos 11-12). (**F**): From these images, the number of protruding cells was quantified (n > 38 fields of view; 3 independent experiments; randomization test).

### MYO10 contributes to ECM assembly *in vitro* and promotes protrusions at the tumor edges *in vivo*

Intrigued by the putative functional role of MYO10-dependent protrusions in ECM assembly, we turned to live *ex vivo* imaging of ECM-embedded tumors. Day 25 xenografts were dissected, embedded in a collagen gel, and imaged at high resolution. With lifeact-mRFP, we observed cells at the border of *in situ* tumor acini extending protrusions that interacted with the surrounding ECM (**Fig. 7C-D**, and **Video 10**). Notably, while very frequent in shCTRL cells, these protrusions were primarily absent in shMYO10 xenografts (**Fig. 7E-F**, and **Video 11-12**). These data indicate that MYO10 contributes to the generation of ECM-probing protrusions at the tumor border, reminiscent of the protruding filopodia detected in the 3D spheroids (**Fig. 7A-B**). While these protrusions are much bigger than individual filopodia, we anticipate that these structures are linked to the ability of the DCIS xenografts to self-assemble the tumor surrounding BM, which is defective in the shMYO10 tumors lacking these dynamic protrusions. Altogether, our data indicate that MYO10 contributes to the protrusive activity of DCIS.com cells *in vivo* in tumor xenografts. These protrusions are essential for proper BM assembly, limiting the DCIS-to-IBC transition. In contrast, silencing of MYO10 correlates with defects in BM assembly *in vivo* and *in vitro* facilitating a transition from DCIS to IBC.

## Discussion

An increasing number of studies indicate a pro-invasive role for MYO10-induced filopodia in aggressive human cancers ranging from lung cancer to melanoma (Arjonen et al., 2014; Cao et al., 2014; Summerbell et al., 2020; Tokuo et al., 2018). Here we set out to study the role of MYO10 at an earlier stage, the transition of in situ tumors to invasive carcinoma. We report that MYO10 expression is increased in patient DCIS lesions which initially led us to hypothesize that MYO10 would facilitate the transition to IBC. Surprisingly, however, the loss of MYO10 expression accelerated rather than protected against invasive progression in a DCIS xenograft model. MYO10-depleted tumors showed defective BM barrier organization at the tumor border and increased dispersal of basal-like cells to the surrounding stroma. Carcinoma cells were identified as the major source of the BM components in vivo and MYO10 depletion induced rather than reduced ECM expression, suggesting defective BM assembly by MYO10-depleted cells in tissue. Concordant with this, functional MYO10 filopodia were found to be essential for BM assembly by 3D tumor spheroids in vitro. Taken together, our observations support a model in which MYO10 plays a dual role in breast cancer progression. In early-stage disease, MYO10 contributes to BM barrier assembly confining DCIS-like tumors in vivo. However, in IBC transitioned tumors, MYO10 filopodia are pro-invasive and contribute to metastasis.

Using high-resolution fixed and live-cell imaging of 3D spheroids, we observed that ECM molecules are recruited by filopodia and deposited at their base. This is consistent with a growing body of evidence of filopodia-like protrusions directly contributing to ECM remodeling in 3D and *in vivo* (Malandrino et al., 2019; Sato et al., 2017; Summerbell et al., 2020) and a report of invading leader cells in collectively migrating lung cancer cells remodeling FN in the shafts of MYO10-positive filopodia (Summerbell et al., 2020). While the precise mechanism(s) by which filopodia remodel ECM remains to be determined, filopodia can assemble adhesive structures capable of interacting with different ECM molecules (Albuschies and Vogel, 2013; Jacquemet et al., 2019; Miihkinen et al., 2021). Filopodia can also exert forces on the underlying substrate, contributing to the remodeling process (Bornschlögl et al., 2013; Brockman et al., 2020; Cojoc et al., 2007; Leijnse et al., 2015). While such a mechanism can easily explain how filopodia-like protrusion can remodel FN (Sato et al., 2017; Summerbell et al., 2020), a process known to require mechanical input from cells (Singh et al., 2010), it is more surprising that filopodia also contribute to the remodeling of BMs as these structures are typically thought to self-assemble (Jayadev and Sherwood, 2017). Our 3D spheroid data indicate filopodia-ECM interaction as an essential mechanism in filopodia-mediated BM assembly. However, further studies are necessary to uncover how filopodia protrusions promote BM assembly.

To invade the surrounding tissues, tumor cells must cross the BM. In this context, BMs are typically viewed as stable barriers that inhibit cancer dissemination. However, recent evidence also suggests that BMs can be very dynamic structures that undergo fast and constant remodeling (Keeley et al., 2020). While, to the best of our knowledge, BM turnover has yet to be observed in tumors, the results presented here would support a model whereby cancer cells contribute to both the production and the assembly of the ECM. Previous work, using proteomic studies, concluded that a large fraction of the ECM in the breast tumor stroma is produced by the cancer cells themselves (Kozma et al., 2021; Naba et al., 2014; Sflomos et al., 2021). Interestingly, tumors lacking MYO10 not only have deficient BM but also have higher ECM gene expression. Therefore, it is tempting to speculate that compensatory mechanisms favor ECM production when ECM assembly is deficient.

DCIS is not life-threatening but becomes potentially lethal in patients who progress to IBC. Therefore, understanding the molecular mechanisms of DCIS-to-IBC transition is an important clinical question. A recent study performed comparative profiling of DCIS epithelium and its microenvironment in samples from DCIS patients with or without subsequent invasive relapse (Risom et al., 2022). Their data indicated that patients with increased stromal cell infiltration, desmoplasia, and ECM remodeling were less likely to relapse to invasive disease (Risom et al., 2022). This study brings forward the intriguing notion of increased stromal desmoplasia correlating with a good prognosis in DCIS, contrary to its widely accepted role in tissue stiffening and cancer progression in IBC (Piersma et al., 2020; Schedin and Keely, 2011).

MYO10 is frequently overexpressed in breast cancer, where its expression correlates with poor prognosis and mutant p53 expression (Arjonen et al., 2014; Cao et al., 2014). MYO10 silencing decreases cancer cell invasion, and metastases of aggressive breast cancer cells *in vitro* and *in vivo* (Arjonen et al., 2014), and MYO10 contributes to the invasion of other aggressive cancers, including melanoma and glioblastoma (Kenchappa et al., 2020; Tokuo et al., 2018). In line with these previous studies, we found that targeting MYO10 expression leads to reduced cell migration *in vitro* and *in vivo*. However, our results also implicate MYO10 plays a dual role in breast cancer progression with a protective, BM supportive function in early-stage disease. While we could not address the metastatic progression in the DCIS xenograft model tested here, defects in BM assembly may accelerate metastasis in other contexts. Therefore, the possible anti- and pro-invasive functions of MYO10 should be carefully considered when therapeutic targeting of this protein is attempted.

## Materials and methods

### Cell lines

MCF10 DCIS.com (DCIS.com) lifeact-RFP cells (Riedl et al., 2008) were cultured in a 1:1 mix of DMEM (Merck) and F12 (Merck) supplemented with 5% horse serum (GIBCO BRL, Cat Number: 16050-122), 20 ng/ml human EGF (Merck, Cat Number: E9644), 0.5 mg/ml hydrocortisone (Merck, Cat Number: H0888-1G), 100 ng/ml cholera toxin (Merck, Cat Number: C8052-1MG), 10 μg/ml insulin (Merck, Cat Number: I9278-5ML), and 1% (vol/vol) penicillin/streptomycin (Merck, Cat Number: P0781- 100ML). The DCIS.com lifeact-RFP cells were generated using lentiviruses, produced using pCDH-lifeAct mRFP, psPAX2, and pMD2.G constructs, as described previously (Jacquemet et al., 2017). The DCIS.com lifeact-RFP shCTRL #s, shMYO10 #3 and shMYO10 #4 cell lines were generated using lentiviruses particles containing a non-target control shRNA (Merck, Cat Number: SHC016V-1EA) or shRNA targeting human MYO10 respectively (shMYO10 #3, TRCN0000123087; shMYO10 #4, TRCN0000123088). Transduced cells were selected using normal media supplemented with 2 µg.ml^-1^ of puromycin. DCIS.com lifeact-RFP shCTRL and shMYO10 lines were generated from single-cell clones obtained from the DCIS.com lifeact-RFP shMYO10 #3 and shMYO10 #4 cell lines. Four single-cell clones with normal MYO10 levels were pooled to create the shCTRL line, and four single-cell clones with very low MYO10 levels were pooled to create the shMYO10 line. The DCIS.com lifeact-RFP shCTRL and shMYO10 GFP lines were generated using lentivirus particles containing GFP. Positive cells were sorted using a BD FACSaria II cell sorter (Becton Dickinson) with a gating strategy to obtain medium expression.

### Generation of tumor xenografts

For xenografts, 1×10^5^ DCIS.com cells were resuspended in 100 µl of a mixture of 50% Matrigel (diluted in PBS) before being injected subcutaneously in the flank or orthotopically in the abdominal mammary gland of 6-7 -week-old virgin female NOD.scid mice (Envigo). Tumor growth was measured with a caliper 1-2 times per week. Mice were sacrificed 10 or 25 days post-injection (as indicated), and the tumors were dissected. For detecting tumor cell proliferation, BrdU Labelling Reagent (Life Technologies) was injected intraperitoneally according to the manufacturer’s instructions (10 µl /g of mouse weight) 2 hours before sacrifice. The National Animal Experiment Board authorized all animal studies, and per The Finnish Act on Animal Experimentation (Animal license number ESAVI-9339-04.10.07-2016; Netherlands Cancer Institute NVWA license number 30100, Project number AVD301002015125).

### *Ex vivo* imaging of tumor xenografts

To perform ex-vivo imaging, DCIS-like xenografts were dissected 25 days post-inoculation, incubated with fibrillar collagen probe CNA35-GFP (when indicated, produced in-house, (Aper et al., 2014)), deposited in a glass-bottom dish (coverslip No. 0; MatTek), and embedded in a collagen-I gel (Advanced BioMatrix, Cat Number: 5074). The gel was then allowed to polymerize at 37 C for 15 min, and the DCIS.com culture medium was added on top. Xenografts were then imaged live using a spinning-disk confocal microscope (40X objective, imaging starting less than 1 h post dissection). Images were processed using Fiji. 3D visualizations were performed using Imaris (Oxford Instruments) and Arivis Vision4D (Arivis).

### Surgical procedures and intravital imaging

Tumor-bearing mice were anesthetized with a 1.5%-2% isoflurane/air mixture 25-35 days post tumor inoculation. To visualize the subcutaneous tumors, a skin flap surgery was performed. The area around the tumor was shaved, disinfected, and a vertical midline incision was made through the skin, followed by two horizontal incisions anterior and posterior of the tumor area. The skin was detached from the underlying tissues and peritoneum by blunt dissection/gentle pulling with a curved instrument. The mouse was transferred to a custom-made imaging box connected to an isoflurane vaporizer. The mouse was placed on top of a metal inlay with a rectangular opening covered with a coverglass. The skin flap was opened, and the tumor area was placed on the coverglass. A sterile gauze soaked in preheated PBS was placed on top of the skin flap to maintain hydration, and parafilm was used to cover the skin flap and create a humidified chamber. To visualize the orthotopic tumors, implantation of an optical imaging window was carried out as described by (Messal et al., 2021). In short, the tumor area was shaved and disinfected, and a 10-15mm incision was made above the tumor. The skin was loosened from the tumor tissue by blunt dissection, and a non-absorbable silk suture was placed in loops around the incision. A sterilized titanium mammary imaging window with a fixed glass coverslip was inserted and secured in the skin with the purse-string suture. After window implantation, the mouse was transferred to a custom-designed imaging box on top of an inlay designed with a hole to secure the imaging window. During time-lapse imaging, the mouse received nutrition through a subcutaneously placed flexible needle (100 µl/hr, Nutriflex (R) special 70/240). Intravital imaging was conducted with an inverted Leica SP8 DIVE microscope (Leica Microsystems) equipped with four tunable hybrid detectors, a MaiTai eHP DeepSee laser (Spectra-Physics), and an InSight X3 laser (Spectra-Physics). For image acquisition, Leica Application Suite X (LAS X) was used.

All images were collected at 12 bit and acquired with a 25 × water immersion objective with a free working distance of 2.40 mm (HC FLUOTAR L 25x/0.95 W VISIR 0.17). GFP and mRFP were excited with 925 nm and 960 nm and detected between 490-550 nm and 580-650 nm, respectively. The second-harmonic generation signal was collected to visualize Collagen I. Whole-tumor areas were imaged by 3D tiles can imaging with a z-step size of 6µm. Timelapse imaging of regions of interest (XYZT) was performed at a 5- or 20-minute time interval for up to 12 hours. Imaged regions were stitched over time using Leica LASX software, and XYZ-drift corrections were performed using Huygens Object Stabilizer software (Scientific Volume Imaging). 3D renderings displayed in Video 4 and 5 were created using the LAS X 3D Visualization module. Motile cells were manually tracked using Imaris software (version 9.0, Oxford Instruments). The mean track speed and persistence were quantified along with the number of invasive (protruding/motile) GFP+ cells per FOV. Image sequences with high cell blebbing (apoptosis due to limited blood supply) were excluded.

### Human tissue samples

Human breast tissue samples were obtained by breast cancer surgery at the Department of Plastic and General Surgery at Turku University Hospital (Turku, Finland) with approval from the Ethics Committee of the Hospital District of Southwestern Finland and written consent from the patients (§279, 9/2001). Human normal breast and breast cancer tissues were collected at Turku University Hospital (Ethical permit 23/1801/2018). Paired samples from breast tumors and surrounding peritumoral or contralateral normal breast tissues of 5 breast cancer patients were excised and examined by a clinical pathologist and, subsequently, processed to formalin-fixed paraffin-embedded (FFPE) tissue sections with standard protocols.

### Antibodies and other reagents

Antibodies used in this studies were anti-Collagen IV (Abcam, Cat Number: ab19808), anti-Fibronectin (FN, Merck, Cat Number: F3648), anti-MYO10 (Novus Biologicals, Cat Number: 22430002), anti-GFP (Abcam, Cat Number: Ab290), anti-alpha-smooth muscle actin (αSMA, clone 1A4, Merck, Cat Number: A2547), anti-Slug (clone C19G7, Cell Signalling Technology, Cat Number: 9585), anti-Vimentin (Clone V9, Santa Cruz, Cat Number: sc-6260), anti-Cleaved caspase-3 (Asp175, clone 5A1E, Cell Signalling Technology Cat Number: 9664), anti-BrdU (clone BU1/75 ICR1, Santa Cruz, Cat Number: sc-56258). The RFP-Booster Atto594 was provided by Chromotek (Cat Number: rb2AF568). Sir-DNA (SiR-Hoechst) (Lukinavičius et al., 2015) was provided by Tetu-bio (Cat Number: SC007). Growth factor reduced Matrigel was purchased from BD Biosciences (Cat Number: 354230). PureCol EZ Gel (fibrillar collagen I, concentration 5 mg/ml) was provided by Advanced BioMatrix (Cat Number: 5074). FITC-collagen was provided by Merk (type I collagen from bovine skin, Cat Number: C4361).

### Histology and immunohistochemistry

Formalin-fixed paraffin-embedded mouse xenograft tissues were sectioned and H&E-labelled with standard procedures. Xenograft histology was scored blindly (*in situ* tumor/ *in situ* tumor with disorganized areas or partial invasion/ tumor with invasion). Immunohistochemistry was performed with standard protocols on deparaffinized sections after heat mediated antigen retrieval in Universal buffer (Retriever 2100, Aptum Bio) with antibodies against KRT8 (clone Troma I, Hybridoma Bank, 1:2000), alpha-smooth muscle actin (αSMA, clone 1A4, Merck, 1:1000), Col IV (ab19808, Abcam, 1:400), FN (F3648, Merck, 1:400), Slug (clone (C19G7, Cell Signalling Technology, 1:100), Vimentin (Clone V9, Santa Cruz, 1:200), Cleaved caspase-3 (Asp175, clone 5A1E, Cell Signalling Technology, 1:500), human mitochondria (clone 113-1, Merck Millipore, 1:100), and BrdU (clone BU1/75 ICR1, Santa Cruz, 1:300). All samples were stained with DAPI (4′,6-diamidino-2-phenylindole, dihydrochloride; Life Technologies), mounted in Mowiol containing DABCO® (Merck) antifade reagent, and imaged with spinning-disk microscopy.

The percentage of GFP-positive cells at the edges of tumor acini was analyzed using Fiji (Schindelin et al., 2012). For each set of samples, four images were acquired: cancer cell marker (human mitochondria), ECM marker (Collagen IV), nuclei (DAPI), and GFP. The edge of tumor acini and its coordinates were first defined using the cancer cell and ECM markers. Then each cell was identified using the DAPI label, and its distance to the closest edge of tumor acini was calculated with R software. Using this information, cells were classified as edge cells (< than 10 µm distance) or not edge cells (> than 10 µm distance). Finally, the GFP channel was used to separate the GFP-positive cells from GFP-negative and quantify the percentage of GFP-positive cells at the edge of tumor acini.

### RNA *in situ* hybridization

RNA *in situ* hybridization was performed on human FFPE breast tissue sections with RNAscope® 2.5 HD Detection kit (BROWN, cat no. 322300) (Advanced Cell Diagnostics) based on manufacturer’s instructions using a probe targeting the region 1262–2318 in MYO10 mRNA (RNAscope® Probe - Hs-MYO10-full, cat no. 440691). For negative and positive controls RNAscope® Negative Control Probe – DapB (cat no. 310043) and RNAscope® Positive Control Probe - Hs-PPIB (cat no. 313901) were used, respectively (Advanced Cell Diagnostics). Nuclei were labeled with hematoxylin, and samples were mounted in DPX new (Merck). Samples were imaged with a Panoramic Slide Scanner (3DHistech).

### Quantitative RT-PCR

Total RNA extracted using the NucleoSpin RNA Kit (Macherey-Nagel) was reverse transcribed into cDNA using the high-capacity cDNA reverse transcription kit (Applied Biosystems) according to the manufacturer’s instructions. The RT-PCR reactions were performed using pre-designed single tube TaqMan gene expression assays (GAPDH: Hs03929097_g1) and were analyzed with the 7900HT fast RT-PCR System (Applied Biosystems). Data were studied using RQ Manager Software (Applied Biosystems).

### Western blotting

Protein extracts were separated under denaturing conditions by SDS–PAGE and transferred to nitrocellulose membranes. Membranes were blocked for one hour at room temperature with a blocking buffer (LI-COR Biosciences) and then incubated overnight at 4 °C with the appropriate primary antibody diluted in the blocking buffer. Membranes were washed with PBS and then incubated with the appropriate fluorophore-conjugated secondary antibody diluted 1:5,000 in a blocking buffer for 30 min. Membranes were washed in the dark and then scanned using an Odyssey infrared imaging system (LI-COR Biosciences).

### Light microscopy

The spinning-disk confocal microscope used was a Marianas spinning-disk imaging system with a Yokogawa CSU-W1 scanning unit on an inverted Zeiss Axio Observer Z1 microscope controlled by SlideBook 6 (Intelligent Imaging Innovations, Inc.). Images were acquired using either an Orca Flash 4 sCMOS camera (chip size 2,048 × 2,048; Hamamatsu Photonics) or an Evolve 512 EMCCD camera (chip size 512 × 512; Photometrics). Objectives used were a 20X air objective (NA 0.8, Plan-Apochromat, Zeiss), a 40x water (NA 1.1, LD C-Apochromat, Zeiss), a 63× oil (NA 1.4, Plan-Apochromat, M27 with DIC III Prism, Zeiss) and a 100x oil (NA 1.4 oil, Plan-Apochromat, M27) objective.

The confocal microscope used was a laser scanning confocal microscope LSM880 (Zeiss) equipped with an Airyscan detector (Carl Zeiss) and a 40x oil (NA 1.4) objective. The microscope was controlled using Zen Black (2.3), and the Airyscan was used in standard super-resolution mode.

### Electron microscopy

The samples were fixed with 5% glutaraldehyde in s-collidine buffer, postfixed with 1% OsO4 containing 1.5% potassium ferrocyanide, dehydrated with ethanol, and embedded in 45359 Fluka Epoxy Embedding Medium kit. Thin sections were cut using an ultramicrotome to a thickness of 70 nm. The sections were stained using uranyl acetate and lead citrate. The sections were examined using a JEOL JEM-1400 Plus transmission electron microscope operated at 80 kV acceleration voltage.

### Proliferation assay

To monitor cell proliferation *in vitro*, cells were plated at low density in a well of a six-well plate and imaged using a live-cell microscopy incubator (IncuCyte ZOOM). Growth rates were calculated using the confluency method within the IncuCyte ZOOM software.

### Circular invasion assays

Cells were plated in one well of a two-well culture-insert (Ibidi, Cat Number: 80209) pre-inserted within a well of a μ-Slide 8 well (Ibidi, Cat Number: 80807). After 24 h, the culture-insert was removed, and a fibrillar collagen gel (PureCol EZ Gel) was cast. The gel was allowed to polymerize for 30 min at 37°C before normal media was added on top. Cells were left to invade for two days before fixation or live imaging.

To analyze filopodia properties, fixed samples were stained with phalloidin-488 and imaged using a spinning-disk confocal microscope (100x objective). Filopodia density and length were then automatically analyzed using the FiloQuant implemented in Fiji (Jacquemet et al., 2017; Schindelin et al., 2012).

To analyze the effect of MYO10 silencing on cell migration, shCTRL, and shMYO10 cells were incubated for 2 h with 0.5 μM SiR-DNA (SiR-Hoechst, Tetu-bio, Cat Number: SC007) before being imaged live for 14 h using a spinning-disk confocal microscope (20x objective, 1 picture every 10 min). Nuclei were automatically detected using the deep learning algorithm StarDist implemented in the ZeroCostDL4Mic platform and tracked using TrackMate (von Chamier et al., 2021; Ershov et al., 2021; Fazeli et al., 2020; Schmidt et al., 2018; Tinevez et al., 2017). This custom StarDist model was trained for 100 epochs on 72 paired image patches (image dimensions: 1024×1024, patch size: 1024×1024) with a batch size of 2 and a mae loss function, using the StarDist 2D ZeroCostDL4Mic notebook (v1.12.2). The StarDist “Versatile fluorescent nuclei” model was used as a training starting point. The training was accelerated using a Tesla P100. This model generated excellent segmentation results on our test dataset (average Intersection over union > 0.96; average F1 score > 0.96) (Laine et al., 2021). The StarDist model and the training dataset used are available for download on Zenodo (Jacquemet, 2020). Cell tracks were further analyzed using the Motility Lab website (http://www.motilitylab.net/, (Wortel et al., 2019)).

To analyze the effect of MYO10 silencing on cell protrusions, shCTRL and shMYO10 cells were imaged live for a few hours using a spinning-disk confocal microscope (100x objective, 1 picture every 3 min). Images were then processed using Fiji (Schindelin et al., 2012).

To perform migration competition assays, GFP positive and negative DCIS.com cell lines were mixed with a 50/50% ratio before being plated in a circular invasion assay. Cells were imaged live for 16 h using a spinning-disk confocal microscope (20x objective, 1 picture every 10 min). For each time point, the migration edges and the GFP-positive cells were automatically segmented and the percentage of the leading edge covered by the GFP-positive cell was then calculated.

### ECM remodeling assay

To form spheroids, DCIS.com cells were seeded as single cells, in standard growth media, at very low density (3,000 cells per well) on growth factor reduced (GFR) Matrigel-coated glass-bottom dishes (coverslip No. 0, MatTek; used for live experiments) or similarly coated glass coverslips (used for fixed samples). After 12 h, the medium was replaced by a standard growth medium supplemented with 2% (vol/vol) GFR Matrigel and 10 µg/ml of a fluorescently labeled ECM molecule (HiLyte Fluor 488-labeled fibronectin, Cytoskeleton, Cat Number FNR02; HiLyte Fluor 488-labeled laminin-111, Cytoskeleton, Cat Number LMN02). After three days, spheroids were imaged live or fixed with 4% PFA for 10 min at room temperature, washed with PBS, and mounted using Mowiol-DABCO.To quantify the amount of ECM recruitment and remodeling occurring at the surface of control and MYO10-depleted DCIS.com spheroids, 3D stacks were acquired using a spinning-disk confocal microscope (step size 0.5 μm). SUM projections were performed, and the integrated intensity was quantified for each spheroid using Fiji (Schindelin et al., 2012).

The impact of different filopodia- and actomyosin-targeting treatments on the ECM remodeling by DCIS.com spheroids was investigated by seeding the cells on GFR Matrigel-coated coverslips as described above. After 12 h, the medium was replaced with a growth medium supplemented with 2% GFR Matrigel, but no fluorescently labeled ECM. The spheroids were grown for two days before labeled fibronectin, and either vehicle (DMSO), 10 µM blebbistatin (STEMCELL Technologies, Cat Number 72402), 10 µg/ml of integrin β1-blocking antibody AIIB2 (in-house production), or 10 µM Saracatinib (Selleck Chemicals, Cat Number S1006) were added. The treatment was carried out for 20 hours, after which the spheroids were fixed, processed into samples, and imaged using a spinning-disk confocal microscope. The three middle planes from each stack (i.e., spheroid) were combined using a SUM projection using a custom Fiji macro. Using F-actin as a marker to indicate the spheroid outlines, the mean intensity of exogenous fibronectin in an approximately 2.5 µm wide region around each spheroid was measured.

For live-cell imaging of the spheroids, a spinning-disk confocal microscope was used. Videos were denoised using the deep Learning algorithm Deconoising (Goncharova et al., 2020) implemented within ZeroCostDL4Mic (von Chamier et al., 2021). The DecoNoising models were trained for 200 epochs directly on the images to denoise using a patch size of 80x 80 pixels, a batch size of 4, and a virtual batch size of 20, using the DecoNoising 2D ZeroCostDL4Mic notebook (v1.12). The training was accelerated using a Tesla P100 GPU, and data augmentation was enabled.

### RNA sequencing and data analyses

Tumors were dissected 25 days after inoculation and stored in an RNAlater lysis buffer (Producer). RNA was extracted from tissue (ca <30mg/sample) collected to H2O using the Qiagen RNeasy Plus Mini kit. The quality of the total RNA samples was ensured with Agilent Bioanalyzer 2100. Sample concentration was measured with Nanodrop ND-2000, Thermo Scientific. Total RNA samples were pure, intact and all samples had a similar quality. Bioanalyzer RIN values were > 9.4. The library preparation was started from 100 ng of total RNA.

Library preparation was done according to Illumina TruSeq® Stranded mRNA Sample Preparation Guide (part # 15031047). The high quality of the libraries was confirmed with Agilent Bioanalyzer 2100, and the concentrations of the libraries were quantified with Qubit® Fluorometric Quantitation, Life Technologies. Library quality was excellent, and all samples had similar quality (fragments in the range of 200-700 bp and the average size of the fragments 250-350 bp).

The samples were normalized and pooled for the automated cluster preparation which was carried out with Illumina cBot station. The 8 libraries were pooled in one pool and run in one lane. The samples were sequenced with the Illumina HiSeq 3000 instrument. Paired-end sequencing with 2 × 75 bp read length was used with 8 + 8 bp dual index run. The technical quality of the HiSeq 3000 run was good, and the cluster amount was as expected. Greater than 75% of all bases above Q30 were requested. The typical yields are 260-310 × 106 paired-end or single-end reads per lane on HiSeq 3000, depending on the library type and quality. The base calling was performed using Illumina’s standard bcl2fastq2 software, and automatic adapter trimming was used.

The quality of the sequencing reads was checked using the FastQC tool (v. 0.11.4) (http://www.bioinformatics.babraham.ac.uk/projects/fastqc). The reads were analyzed against both human and mouse references. First, the sequencing reads were separately aligned to human (UCSC hg38) and mouse (UCSC mm10) reference genomes, derived from the Illumina iGenomes resource (https://support.illumina.com/sequencing/sequencing_software/igenome.html), using STAR aligner (v. 2.5.2b) (Dobin et al., 2013). For mouse reference, the reads that also aligned to human reference were removed using XenofilteR (v. 0.99.0) (Kluin et al., 2018). Next, the uniquely aligned reads were associated with RefSeq gene models using Subread (v. 1.5.1) (Liao et al., 2014) for each organism. Normalization and statistical testing were carried out with R (v. 3.4.1) and Bioconductor (v. 3.6) (Gentleman et al., 2004), using edgeR (McCarthy et al., 2012) and Limma packages (Ritchie et al., 2015). In each comparison, genes with mean RPKM expression value below 0.125 in both sample groups were filtered out, and the normalized expression values were voom transformed before statistical testing. An absolute fold-change above two and a false discovery rate (FDR) smaller than 0.01 or 0.05 were required to select the differentially expressed genes.

### Quantification and statistical analysis

Randomization tests were performed using PlotsOfDifferences (Goedhart, 2019). Dot plots were generated using PlotsOfData (Postma and Goedhart, 2019), Volcano Plots were generated using VolcaNoseR (Goedhart and Luijsterburg, 2020), and SuperPlots were generated using SuperPlotsOfData (Goedhart, 2021; Lord et al., 2020). Other statistical tests were performed using GraphPad Prism software. The Student’s t-test (unpaired, two-tailed) was used for normally distributed data. Non-parametric Mann–Whitney U-test was used when two non-normally distributed groups were compared. Fisher’s exact test was used for the analysis of contingency tables. Data representation and n-numbers for each graph are indicated in figure legends.

## Data and code availability

The StarDist model used to track cells automatically is available on Zenodo (Jacquemet, 2020). The RNA sequencing dataset generated here has been added to Gene Expression Omnibus (GEO) database repository (GSE166898). The authors declare that the rest of the data described in this manuscript are also available upon request.

## Acknowledgments

We thank H. Hamidi for providing feedback and editing the manuscript. The Cell Imaging and Cytometry Core facility (Turku Bioscience, University of Turku, Åbo Akademi University, and Biocenter Finland), the Finnish Functional Genomics Centre (Turku Bioscience, University of Turku, Åbo Akademi University, and Biocenter Finland), the Laboratory of Electron Microscopy and Histocore (Institute of Biomedicine, University of Turku), Central Animal Laboratory (University of Turku), and Turku Bioimaging are acknowledged for services and instrumentation and expertise. We thank Ilkka Koskivuo for providing us with clinical breast cancer tissue samples, and Mitro Miihkinen, Artur Padzik, and Petra Laasola for assistance during the experimental work.

This work was supported by the Finnish Cancer Institute (K. Albin Johansson Professorship to J.I.), Academy of Finland Research project grant (325464 to J.I.), Finnish Cancer Organization (grants to J.I. and G.J.), Sigrid Juselius Foundation (grants to J.I., E.P., and G.J.), Academy of Finland research fellowships (323096 to E.P., 338537 to G.J., 312517 to M.G.), the Hospital district of Southwest Finland (11083 to E.P.), the University of Turku Foundation (grants to E.P. and G.J.), the Åbo Akademi University Research Foundation (CoE CellMech; to G.J.), the Drug Discovery and Diagnostics strategic funding to Åbo Akademi University (to G.J.), the Boehringer Ingelheim Foundation (Ph.D. fellowship to C.L.G.J.S.), EMBO postdoctoral fellowship (grant ALTF 1035-2020 to C.L.G.J.S.), Finnish Cultural Foundation (to A.I.), Josef Steiner Cancer Research Foundation (to J.v.R.).

## Author contributions

Conceptualization, E.P., G.J., J.I.; Methodology, E.P., G.J., J.I., C. L.G.J. S.; Formal Analysis, E.P., G.J., C.L.G.J.S., A. I., C. G., S.K., A.L., L.L.E., M. G.; Investigation, E. P., G. J., C. L.G.J. S., A.I., I. P., K. T., M. G., P. B., I.P.; Writing – Original Draft, E.P., G.J., J.I.; Writing – Review and Editing, All authors; Visualization, E.P., G.J., A.I., J.I.; Supervision, E.P., G.J., J.I., L. L. E., J. v. R.; Funding Acquisition, E.P., G.J., J.I..

## Video legends

**Video 1**: shCTRL and shMYO10 MCF10DCIS.COM cells, labeled with Sir-DNA, migrating through a collagen gel were recorded using a spinning-disk confocal microscope (20x objective). Cells were then automatically tracked using StarDist and TrackMate. Raw data and Local tracks are displayed. Colour indicates ID.

**Video 2**: shCTRL and shMYO10 MCF10DCIS.COM cells migrating through a collagen gel were recorded using a spinning-disk confocal microscope (100x objective) to visualize the protrusions generated at the invasive front.

**Video 3**: Various shCTRL and shMYO10 DCIS.com cell lines were mixed in different combinations so that one of the cell lines is always GFP positive, and their migration was recorded live on a spinning-disk confocal microscope (20x). The GFP-positive cells (green) and the invasive edge (magenta) were thresholded using Fiji (bottom panels).

**Video 4:** shCTRL^GFP^ + shMYO10 DCIS.com cells were xenografted in NOD.scid mice in 1:1 ratio. After 33 days, the resulting xenografts were imaged by intravital tile scan imaging. The 3D reconstruction was performed using the LAS X 3D Visualization module.

**Video 5:** shCTRL + shMYO10^GFP^ DCIS.com cells were xenografted in NOD.scid mice in 1:1 ratio. After 27 days, the resulting xenografts were imaged by intravital tile scan imaging. The 3D reconstruction was performed using the LAS X 3D Visualization module.

**Video 6**: shCTRL^GFP^ were injected s.c. with non-GFP shMYO10 cells and imaged with intravital microscopy through a skin flap 34 days post-tumor inoculation. Scale bar: 30 μm, interval 5 min; duration 3 hours 20 min.

**Video 7**: DCIS.com cells were seeded as single cells in Matrigel. They were allowed to form spheroid for three days. Spheroids were then imaged live using a spinning-disk confocal microscope (63X).

**Video 8**: DCIS.com cells were seeded as single cells in Matrigel. They were allowed to form spheroid for three days in the presence of fluorescently labeled fibronectin. Spheroids were then imaged live using a spinning-disk confocal microscope (63X). Images were then denoised using DecoNoising implemented within ZeroCostDL4Mic.

**Video 9**: Lifeact-mRFP-expressing DCIS.com cells were seeded as single cells in Matrigel, allowed to form spheroids for two days, supplemented with fluorescently labeled fibronectin and either 0.1% DMSO or 10 µM Saracatinib, and grown for an additional 20 hours before fixing, mounting, and imaging with a spinning-disk confocal microscope (63x). Full 3D volumes of two representative spheroids are displayed.

**Video 10**: Freshly dissected day 25 DCIS-like xenografts (green) were imaged live ex-vivo in the presence of CNA35-GFP (magenta) using a spinning-disk confocal microscope (40X, ORCA camera). The 3D reconstruction was performed using Arivis Vision4D.

**Video 11**: Freshly dissected day 25 shCTRL DCIS-like xenografts were imaged live ex-vivo using a spinning-disk confocal microscope (40X, ORCA camera). The 3D reconstruction was performed using Arivis Vision4D.

**Video 12**: Freshly dissected day 25 shMYO10 DCIS-like xenografts were imaged live ex-vivo using a spinning-disk confocal microscope (40X, ORCA camera). The 3D reconstruction was performed using Arivis Vision4D.

**Figure S1:**
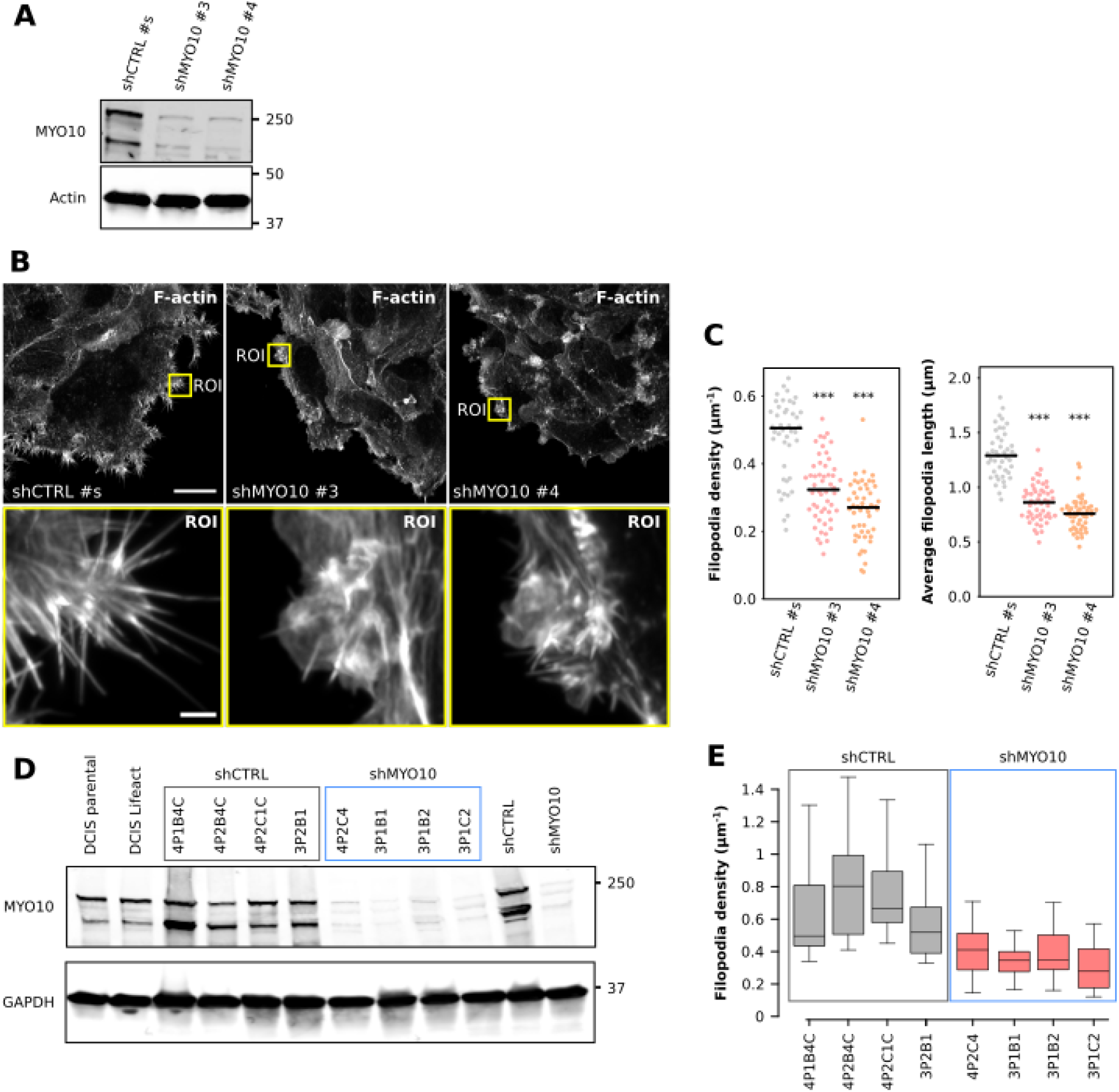
MYO10 modulates filopodia formation. (**A**): DCIS.com cells were infected with lentiviruses containing shRNA targeting MYO10 or CTRL shRNA. After antibiotic selection, cells were lysed, and MYO10 protein levels were analyzed by western blot. (**B-C**): DCIS.com cells generated in (**A**) were left to migrate underneath a collagen gel for 2 d, fixed, stained, and imaged using a spinning-disk confocal microscope. (**B**): Representative fields of view are displayed showing the morphology of shCTRL #s, shMYO10 #3, and shMYO10 #4 cells. Yellow squares highlight ROIs that are magnified. Scale bars: (main) 25 μm; (inset) 2 μm. (**C**): Filopodia density and the average filopodia length were analyzed using FiloQuant. Results from three independent experiments are displayed as dot plots (n > 44 fields of view analyzed per condition; *** p-value < 0.001, randomization test). (**D-E**): Single-cell clones were isolated from the cell lines generated in (**A**), and their MYO10 protein levels were compared to parental DCIS.com cells by western blot. (**D**): Four clones with low MYO10 expression and four clones with high MYO10 expression were mixed to generate the shMYO10 and shCTRL cell lines, respectively. (**E**): Filopodia density for each single cell clone was automatically analyzed using FiloQuant. Results are displayed as box plots (n > 20 fields of view analyzed per condition; two independent experiments). (**F**): DCIS.com cells were injected subcutaneously in NOD.scid mice. The resulting tumors were dissected ten, 17, or 24 d post-injection. Tumor sections were then stained using H&E and imaged. Representative images are displayed. Scale bars: 100 μm.

**Figure S2:**
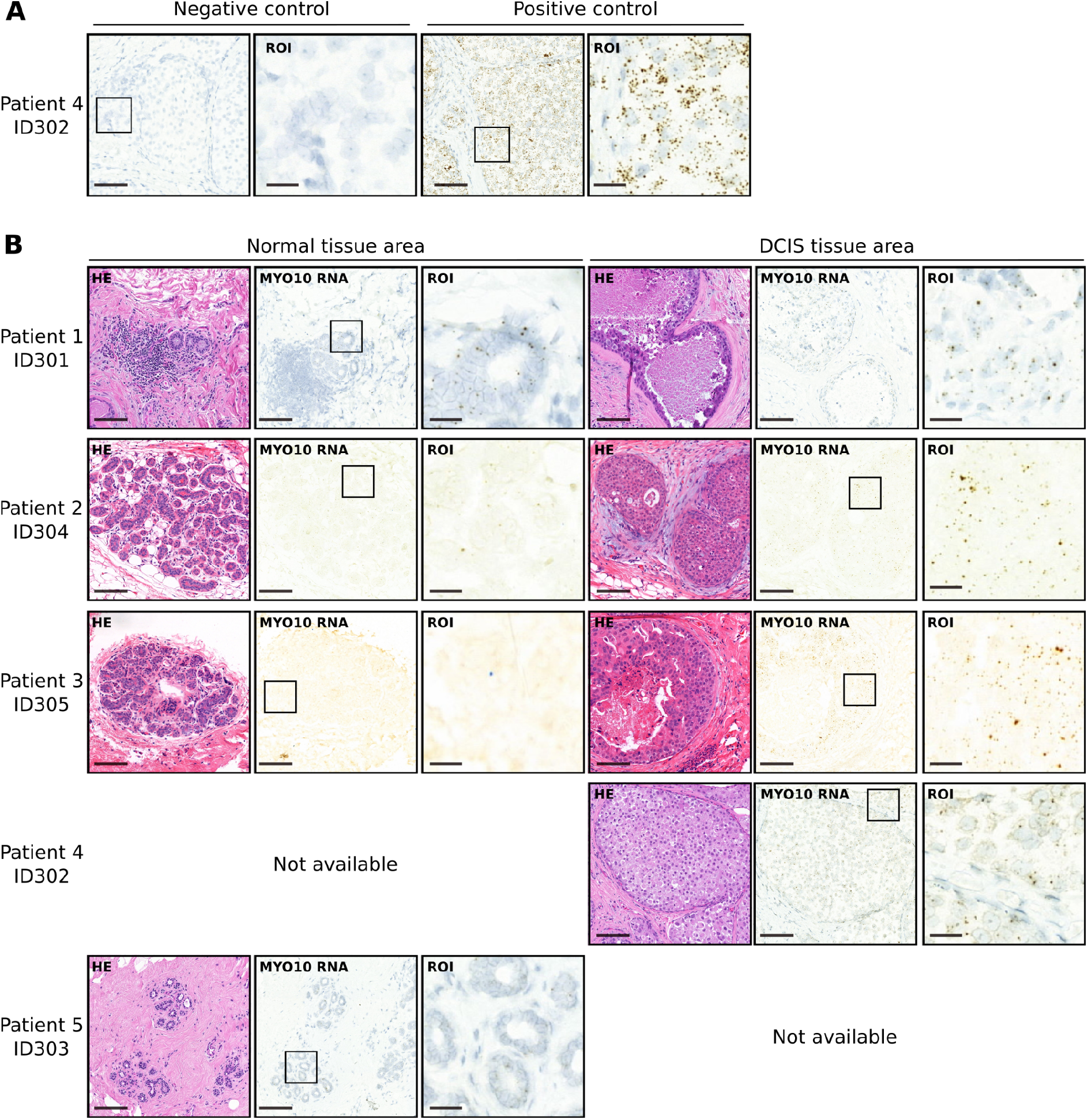
MYO10 mRNA expression in patients undergoing lumpectomy after DCIS diagnosis. (**A**): Representative images of RNA *in situ* labeling with negative (DapB) and positive (Hs-PPIB) control probes in DCIS regions of a human breast sample (nuclei labeled with hematoxylin). RNA can be visualized by the dots visible in the magnified ROI.) Negative and positive controls were included in all independent experiments. (**B**): Representative RNA *in situ* labeling of MYO10 mRNA in normal and DCIS regions of a human breast sample (4 patient samples per condition). MYO10 mRNA can be visualized by the dots visible in the magnified ROI. Patient 4 sample did not contain any areas of normal breast, and patient 5 did not have DCIS despite the initial diagnosis. Elevated MYO10 expression was detected in DCIS areas compared to histologically normal breast tissue. Scale bars: (main) 100 μm; (inset) 20 μm.

**Figure S3:**
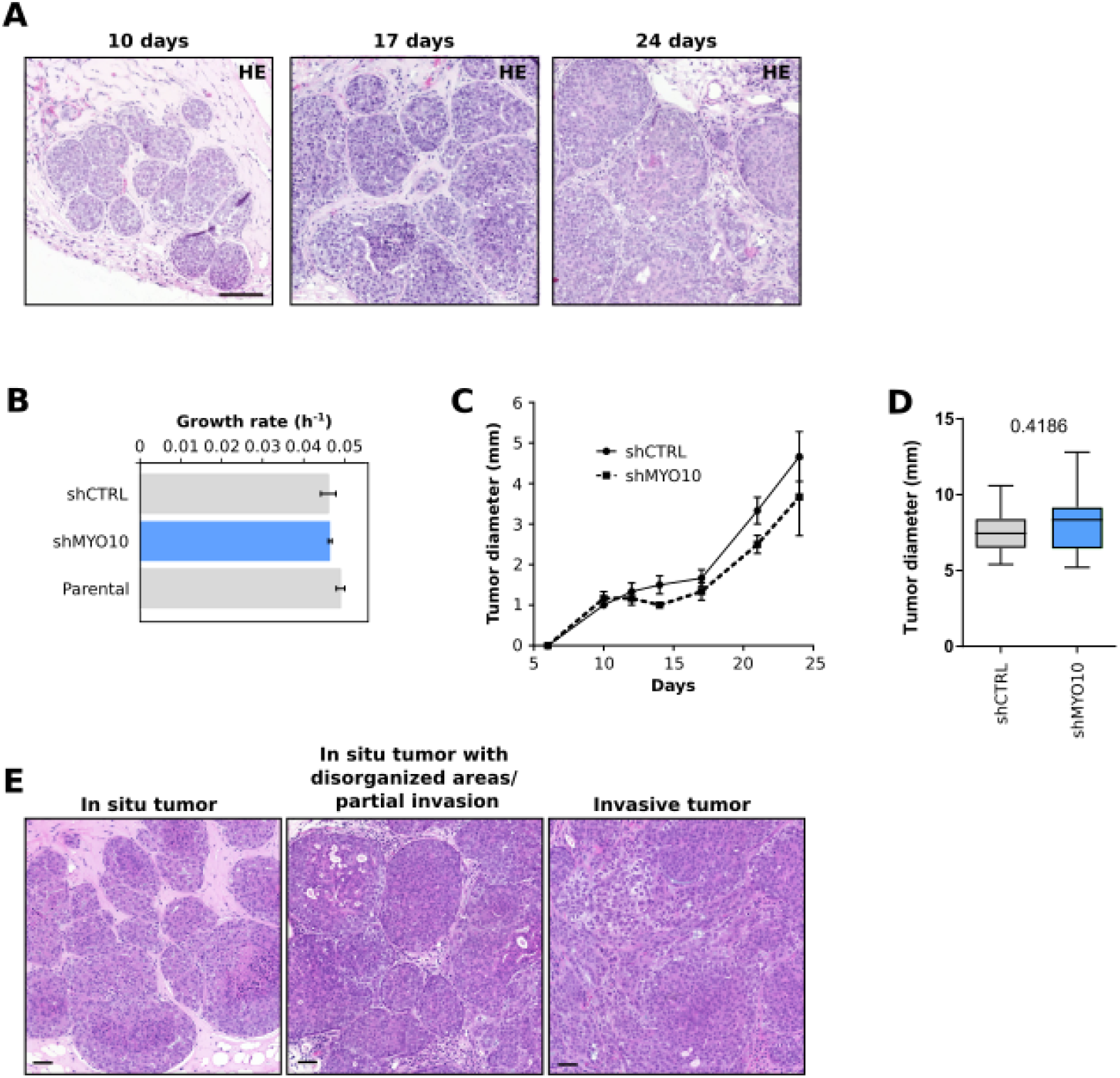
MYO10 is not required for DCIS.com cells proliferation *in vitro* or *in vivo*. (**A**): DCIS.com cells were injected subcutaneously in NOD.scid mice. The resulting tumors were dissected ten, 17, or 24 d post-injection. Tumor sections were then stained using H&E and imaged. Representative images are displayed. Scale bars: 100 μm. (**B**): shCTRL and shMYO10 cells were plated in a six-well plate, and their proliferation rate was recorded using an incubator microscope (three biological repeats, error bars indicate SEM). (**C-D**): shCTRL and shMYO10 cells were injected subcutaneously in immunocompromised mice, and (**B**): the tumor diameter was followed over time (n= 6 tumors). (**C**): At 25 days post-injections, the tumors were dissected and their maximum diameter measured (n=12 tumors, from 3 independent experiments, unpaired t-test). (**E**): Images of H&E stained tumors representing different categories in histological analysis. Scale bars: 50 μm.

**Figure S4:**
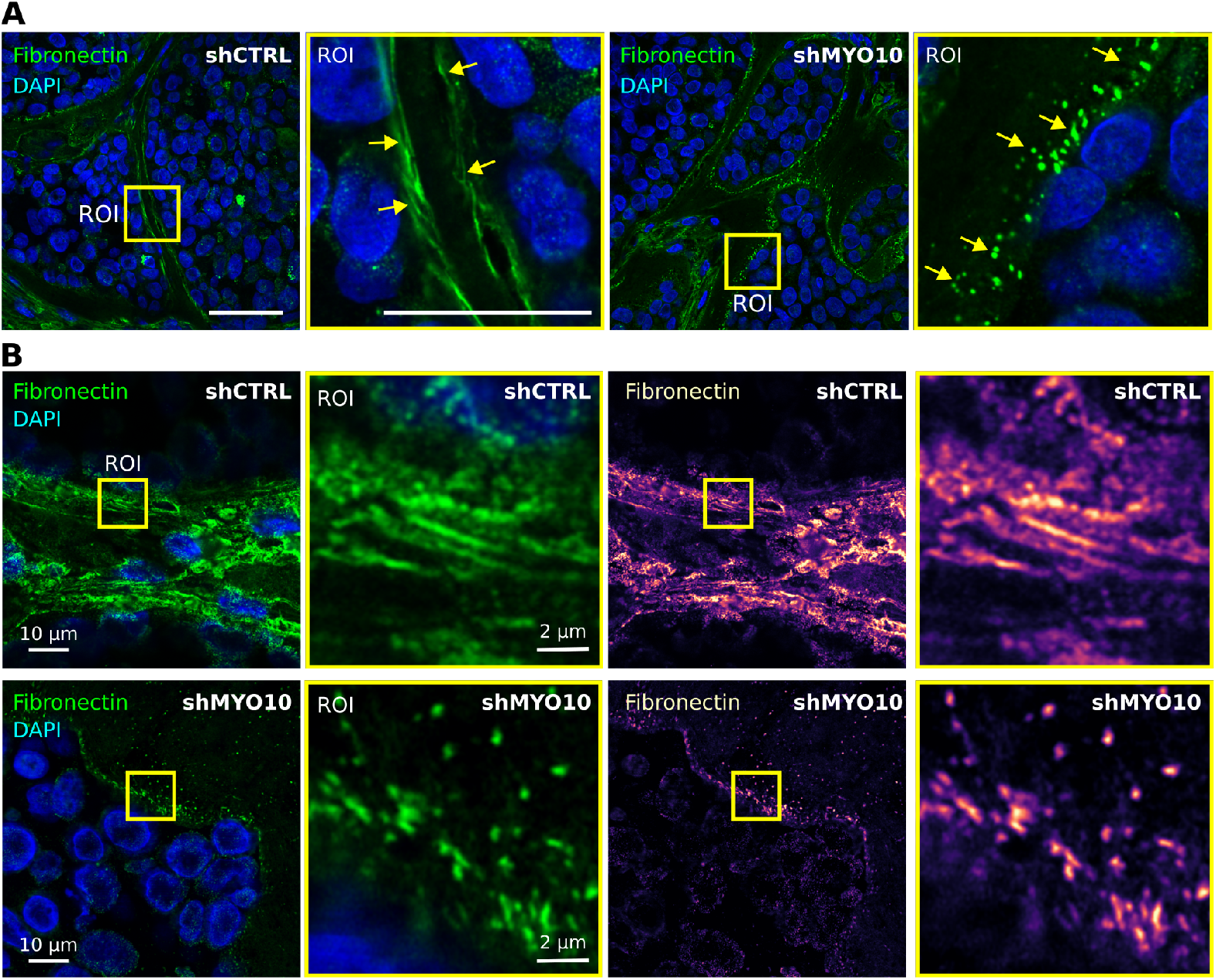
MYO10 modulates fibronectin assembly *in vivo*. (**A-B**): Ten-day-old shCTRL and shMYO10 DCIS-like xenografts were stained for DAPI and fibronectin and imaged using a spinning-disk confocal microscope (**A**, 63x objective) or at high resolution using an Airyscan confocal microscope (**B**). Representative fields of view are displayed. Yellow squares indicate ROIs that are magnified. (**A**): Scale bars: (main) 50 μm; (inset) 25 μm. (**B**): Scale bars: (main) 10 μm; (inset) 2 μm.

**Figure S5:**
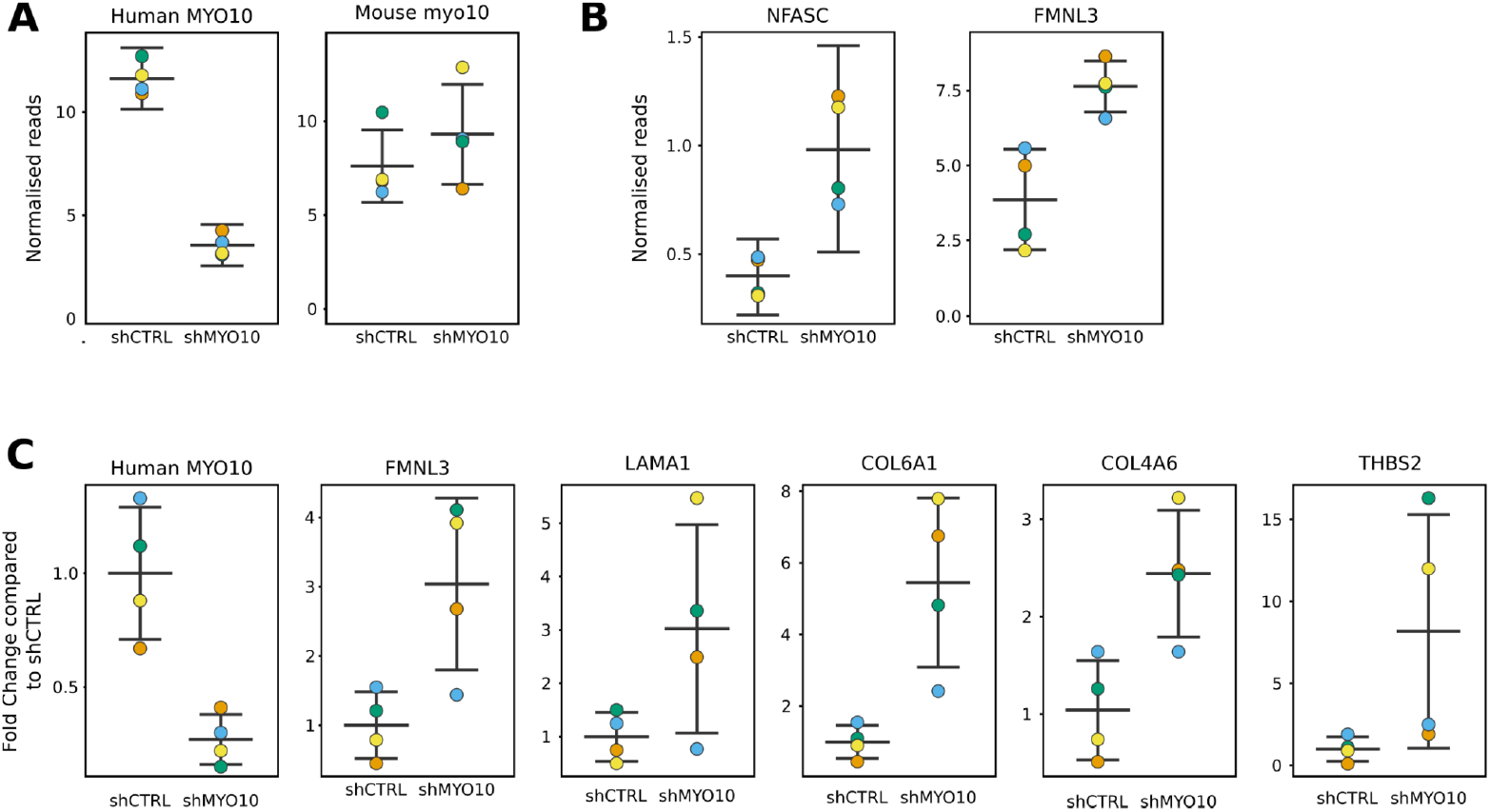
MYO10 silencing drives the expression of filopodia-inducing and ECM genes. (**A-C**): The expression levels of multiple genes (as indicated) were measured in 25-day-old shCTRL and shMYO10 DCIS-like xenografts using RNAseq (**A, B**) or qPCR (**C**), and results are displayed as SuperPlots (Goedhart, 2021; Lord et al., 2020). N = 4 mice per condition.

